# Long-term mavacamten exposure reduces force and sarcomere density in a hiPSC model of hypertrophic cardiomyopathy

**DOI:** 10.1101/2025.10.14.681400

**Authors:** Marianna Langione, Lucrezia Giammarino, Roberto Semeraro, Beatrice Scellini, Sonette Steczina, Valentina Spinelli, Elema Martelli, Irma Della Corte, Camilla Olianti, Alberto Magi, Leonardo Sacconi, Elisabetta Cerbai, Michael Regnier, Iacopo Olivotto, Chiara Tesi, Corrado Poggesi, Raffaele Coppini, Cecilia Ferrantini, J. Manuel Pioner

**Author notes:** **Corresponding authors**: Marianna Langione and J. Manuel Pioner, Department of Biology, University of Florence, Florence Italy **Email:**.

## Abstract

**Background:** Mavacamten, a first-in-class allosteric myosin inhibitor, has demonstrated efficacy and safety in obstructive hypertrophic cardiomyopathy (oHCM), notably reducing symptoms, left ventricular outflow obstruction, and wall thickness over 30 weeks. We recently reported that the *MYBPC3*:c.772G>A variant causes HCM through cMyBP-C haploinsufficiency, leading to accelerated sarcomere kinetics and higher energy consumption in patient myocardium and hiPSC- derived cardiomyocytes (hiPSC-CMs). These effects are counterbalanced by prolonged action potentials and slower Ca²⁺ transients, which preserve twitch duration but may increase arrhythmic risk. Mavacamten may reduce myocardial energetic defects in HCM.

**Objectives:** To investigate the long-term effects of Mavacamten on sarcomere structure, contractility, and transcriptional remodeling using patient-specific and CRISPR-corrected isogenic hiPSC-derived cardiomyocyte models of HCM.

**Methods:** HiPSC-CMs and engineered heart tissues (EHTs) derived from a *MYBPC3:c.772G>A* patient and its CRISPR-corrected line were first exposed to increasing concentrations of Mavacamten to assess acute dose–response relationships and determine IC_50_ values. Based on these data, chronic treatments (0.3– 0.75 μM for 20 days) were performed mechanical, structural, electrophysiological, and transcriptomic adaptations.

**Results:** Acute exposure produced a rapid and fully reversible reduction in active force, while chronic treatment for 20 days induced a sustained decrease in contractility with incomplete recovery after 4 days of washout, indicating a two-phase mechanism of action. Long-term force reduction was paralleled by decreased cell area and sarcomere density, indicating that structural disassembly contributes to sustained functional depression and re-assembly after washout. Electrophysiological analysis confirmed the specific alterations of the *MYBPC3*:c.772G>A mutation previously observed, with no detectable effects following treatment with Mavacamten. In addition, transcriptome analysis was used to study the molecular mechanisms underlying the long-term effect.

**Conclusions:** Mavacamten induces a biphasic, persistent-to-reversible, reduction of sarcomeric force associated with structural remodeling, providing mechanistic insight into its capacity to promote favorable cardiac remodeling in oHCM.

## Introduction

Hypertrophic cardiomyopathy (HCM) is the most common Mendelian heart disease often defined as a disease of the sarcomere, as it is frequently caused by mutations in genes encoding sarcomere proteins (1,2). HCM is commonly associated with myocardial hypercontractility and increased energy cost of sarcomere contraction, defects recently identified as a suitable target for disease-specific pharmacological approaches (3–5). In several HCM-related genetic backgrounds involving myofilament genes, myocardial hypercontractility has been attributed to the increased number of myosin heads available (4). Mavacamten, a first-in-class allosteric inhibitor of myosin ATPase, acts by stabilizing the super relaxed state (SRX) of myosin heads in cardiac muscle (6,7) countering hypercontractility, thus normalizing myocardial energetics in experimental HCM models. In single human ventricular myofibrils, Mavacamten induces acceleration of cross-bridge detachment and inhibition of the ADP-stimulated force developed in the virtual absence of external Ca^2+^ (8). In human ventricular strips Mavacamten maintained the length-dependent activation response even though it reduced myofilament Ca^2+^ sensitivity (9). In mice carrying pathogenic cardiac myosin heavy chain mutations, early administration of Mavacamten prevented the development of hypertrophy, cardiomyocyte disarray, and fibrosis, and mitigated pathological gene expression (3). These findings support its use as a sarcomere-targeting therapy for HCM patients, with the goal of restoring energetic balance at the cellular level.

In phase 2 and 3 clinical studies, Mavacamten consistently demonstrated efficacy and safety in reducing symptoms and left ventricular outflow tract obstruction, improving biomarker profile and reduced exercise capacity, in patients with obstructive HCM (5,10). In addition, Mavacamten has shown disease-modifying properties including reduction in maximum left ventricular (LV) wall thickness and LV mass, compared to placebo in the relatively short-term of 30 weeks (11,12). Notably, these effects are not observed with traditional treatments of obstructive HCM, such as betablockers or disopyramide. Yet, all these improvements in key markers of HCM pathophysiology have been demonstrated over the short term, while the long-term impact of myosin heavy chain inhibition remains unclear. The MAVA-LTE (Mavacamten Long-Term Extension study; NCT03723655) and other longitudinal studies are in the process to better clarify the clinical impact of long-term Mavacamten effect on HCM patients.

We recently described a large and well-characterized population carrying the *MYBPC3:*c*.772G>A* variant (13), which represents a founder effect in the Tuscany region (14). This variant provided the unique opportunity to study the basic mechanisms of *MYBPC3* mutation causing HCM using patient myocardium from surgical samples, as well as induced pluripotent stem cell-derived models (hiPSCs) derived from one of our patients. In the myofibrils of patients and hiPSC-CMs carrying the *MYBPC3:*c*.772G>A* variant, cardiac myosin binding protein C (cMyBP-C) haploinsufficiency causes acceleration of cross bridge kinetics and increased energetic cost of contraction (14,15). Interestingly, electrophysiological changes (slower action potentials and prolonged calcium transients) appeared to counterbalance the faster cross-bridge cycling, ultimately preserving the amplitude and duration of cardiac contraction, at the cost of reduced cardiac electrical stability (development of early and delayed afterdepolarizations) and impaired diastolic function (14,16).

The advantage of using both the HCM patient myocardium and related patient-derived hiPSC line allowed us to better demonstrate the predictive power of patient-specific hiPSC-CMs for modelling inherited cardiomyopathies. The fact that the same changes observed in cardiomyocytes from patient tissue were recapitulated by the *MYBPC3:c.772G>A* hiPSC-CMs and engineered heart tissues (EHTs), suggested that these mechanisms appear early during disease development, likely in the pre- hypertrophic stages of the disease. Thus, these models offer an opportunity for a deeper mechanistic interpretation of the long-term disease-modifying impact of Mavacamten.

In the present study, we therefore leveraged an isogenic pair of hiPSC lines derived from a patient carrying the *MYBPC3*:c.772G>A variant and its CRISPR-corrected counterpart to examine the long- term effects of Mavacamten. We generated cardiomyocytes (hiPSC-CMs) and engineered heart tissues (EHTs) from these lines and assessed structural, functional, and transcriptomic responses to chronic Mavacamten exposure. Our goal was to better understand the molecular and cellular mechanisms underlying its therapeutic action and potential off-target effects across HCM disease stages. Our findings provide a comprehensive insight into the contribution of long-term exposure to Mavacamten to reverse remodelling in HCM.

## Results

### Impact of the MYBPC3:c772G>A mutation on the hiPSC-EHTs function

To investigate the targets and effects of Mavacamten, we employed hiPSC obtained from a previously characterized HCM patient line (ID3) carrying the *MYBPC3:c772G>A* mutation and its CRISPR- Cas9 corrected isogenic control (c.ID3). Differentiated hiPSC-CMs were used to generate the engineered heart tissues (EHTs), as previously reported (14). Approximately 8 days after hiPSC-CM seeding into fibrin scaffold, the EHTs developed spontaneous contractions. At day 60 post differentiation (p.d.) EHTs were manually detached from the pillars and mounted on a force recording apparatus under isometric fixed end conditions under imposed pacing rates. Compared to the CRISPR-Cas9 corrected EHTs (c.ID3), mutant EHTs (ID3) showed no significant differences in contraction amplitude (c.ID3 1.05±0.34 mN/mm²; ID3 1.24±0.22 mN/mm²) (**Figure 1A-B**) and similar time-course of contraction (c.ID3 335.55±22.70 ms; ID3 330.69±26.94 ms) (1.8 mM [Ca^2+^]; 1Hz) (**Figure 1C**). At the same day of maturation, action potential (AP) and Ca^2+^ transients (CaT) recordings were investigated using fluorescent dyes on a confocal microscope. As observed in the **Figure 1D-E**, ID3-EHTs showed a prolonged AP (370.19±10.93 ms; 1 Hz) and slower calcium transients (365.38±55.71ms; 1 Hz) than in controls EHTs (c.ID3) (AP 210.41±48.13 ms; CaT 237.34±48.67 ms; 1 Hz), recapitulating what was observed in cardiomyocytes isolated from native tissue and single hiPSC-CMs (14). EHTs were then exposed in acute condition to Mavacamten to obtain a dose-response curve. Steady-state active tension decreased rapidly to a new steady state (50% of reduction) in both ID3 and c.ID3 (**Figure 2A**). IC_50_ was calculated in acute condition at 1.8 mM [Ca^2+^] was 0.75 µM and 0.47 µM for ID3 and c.ID3 respectively (**Figure 2B-C**). A separate set of experiments was carried out also to verify how quickly Mavacamten can be removed from the medium and reverse the contractile properties of EHTs. A group of untreated ID3-EHTs was measured under isometric fixed-end conditions at the end stage of maturation (d60 p.d.) and rapidly exposed to Mavacamten (0.75 µM) (**Figure 2D**). Consistent with the previous test, the steady-state active tension decreased rapidly to a new steady state (50% reduction). After an immediate washout, steady-state tension returned to the initial level within about one hour (**Figure 2E**), demonstrating the reversibility of the Mavacamten effect after washout.

**Figure 1.**
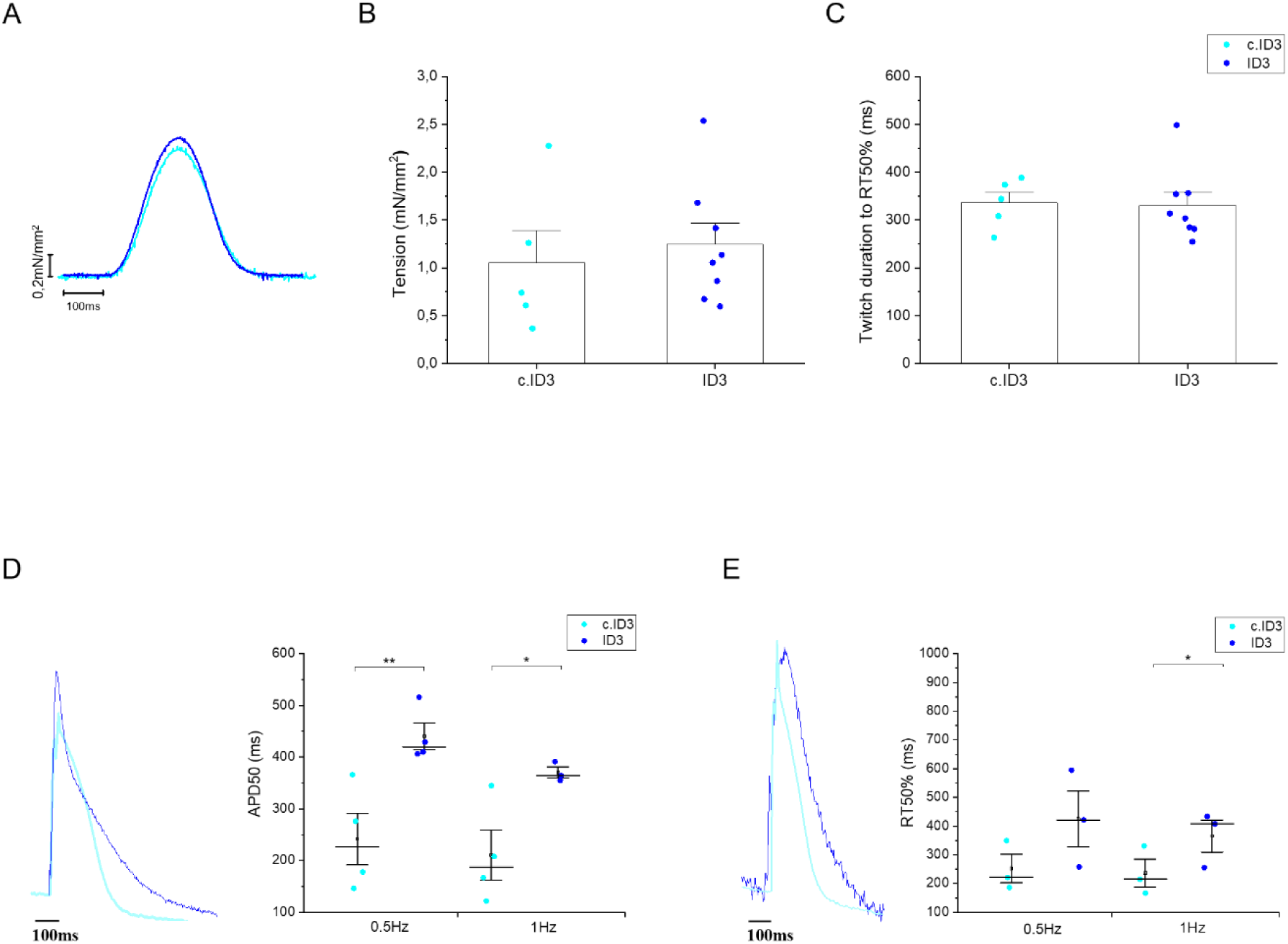
Impact of the MYBPC3:c772G>A mutation on the function of hiPSC-EHTs. (A-B) Active tension and (C) twitch duration at 50% of relaxation of isogenic control (c.ID3) (N=4; n=5) compared with c.772G>A-EHTs (ID3) (N=5; n=8) at 1Hz constant pacing with 1.8 mM of [Ca^2+^]. (D-E) Simultaneous recording of action potential and calcium transients by fluorescent indicators (Cal630 and FluoVolt, respectively). Single mutant hiPSC-EHTs (ID3, AP N=4, n=4; CaT N=2, n=3) were compared with its isogenic control (c.ID3, AP N=3, n=4; CaT N=3, n=3) for action potential at 50% of duration (APD50, ms) and the duration from calcium transient peak to 50% of calcium transient decay (RT50, ms). Error bars ± SEMs. One-way analysis of variance (ANOVA) with a Tukey post-hoc test was used. * p < 0.05 and ** p < 0.01.

**Figure 2.**
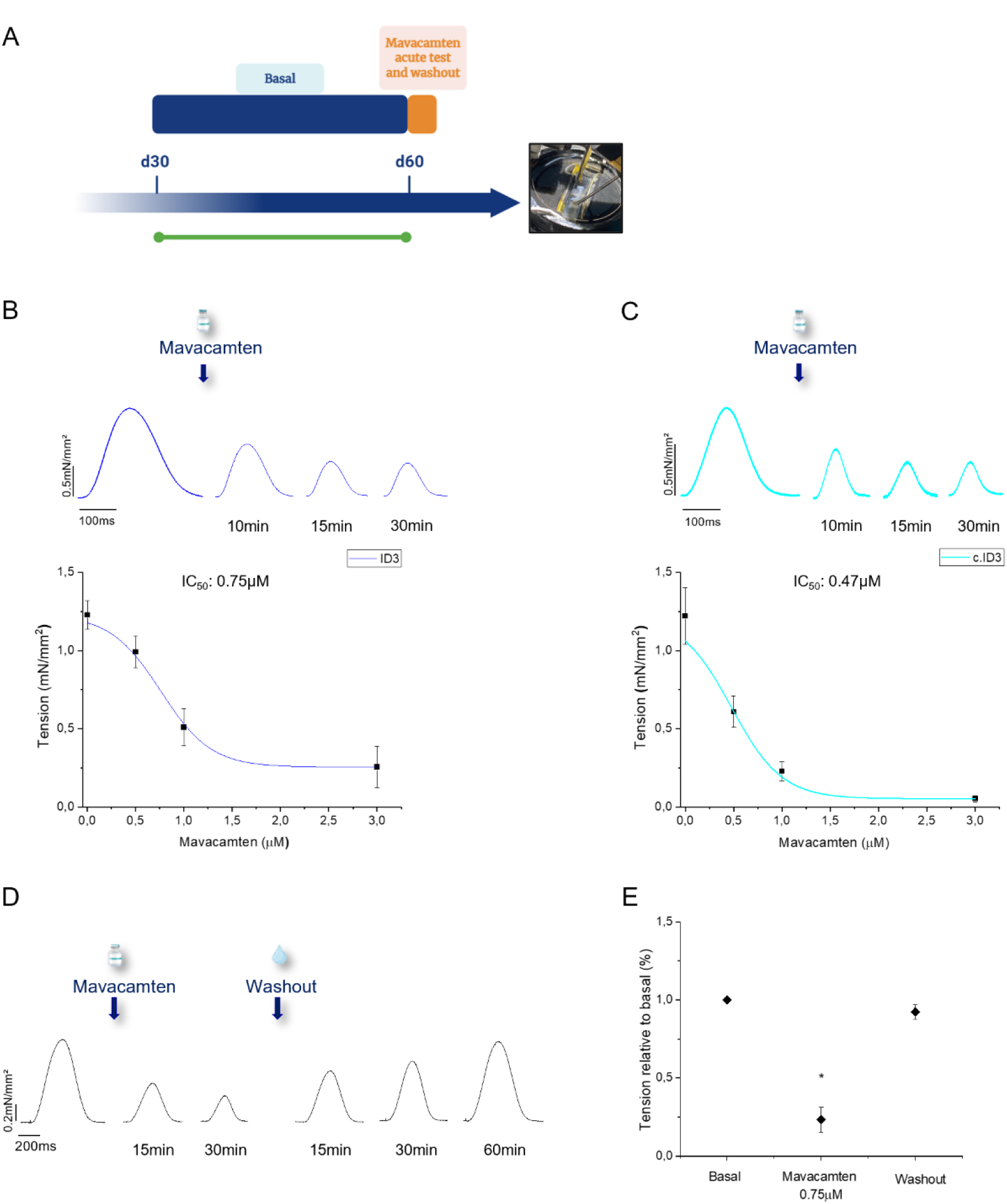
Effect of mavacamten on contractile force under isometric conditions. **(A)** Schematic representation of the acute test with Mavacamten. The scheme was created with BioRender.com. **(B-C)** Representation of the dose-response curves of Mavacamten in HCM- (ID3) (n=4) and isogenic control-EHTs (c.ID3) (n=4). Representative twitches traces **(D)** and **(E)** active tension at 1Hz before, during, and after acute treatment with Mavacamten 0.75µM (N=2, n=2) ID3-EHTs. EHTs were measured under isometric conditions at 37°C in Krebs-Henselheit solution with 1,8mM of [Ca^2+^]. Error bars ± SEMs.

### Chronic effect of Mavacamten in Engineered Heart Tissues

#### Spontaneous auxotonic contractions of EHTs during chronic treatment with Mavacamten

The present study sought to assess any functional changes that may occur after prolonged treatment with Mavacamten. c.ID3- and ID3-EHTs were exposed for 20 days to two different concentrations of Mavacamten, at 0.3 µM, as in other works (17,18), and 0.75 µM (**Figure 3A**), in agreement with the ID3 IC_50_. For the control group a concentration of 0.75 µM DMSO was added to the culture medium. Treatment with Mavacamten was started 10 days after EHT generation. Before and during treatment, the tension generated by spontaneous auxotonic contractions was recorded at specific time points (**Figure 3B-G**, **Video S1**). The untreated c.ID3 and ID3 EHTs exhibited a gradual increase in tension over time in culture, reaching a comparable level (**Figure 3C and 3G**). In contrast, Mavacamten-treated EHTs showed a dose-dependent reduction in spontaneous force: tension was reduced (0.3 µM) or disappeared (0.75 µM) in both ID3 and c.ID3 EHTs (**Video S2-S3**). After 20 days of treatment, Mavacamten was removed from the culture medium (washout), spontaneous active tension was measured in treated EHTs showing tension recovery, suggesting that the effects of Mavacamten are reversible (**Figure 3D**, **3E, 3H and 3I**, **Video S4**).

**Figure 3.**
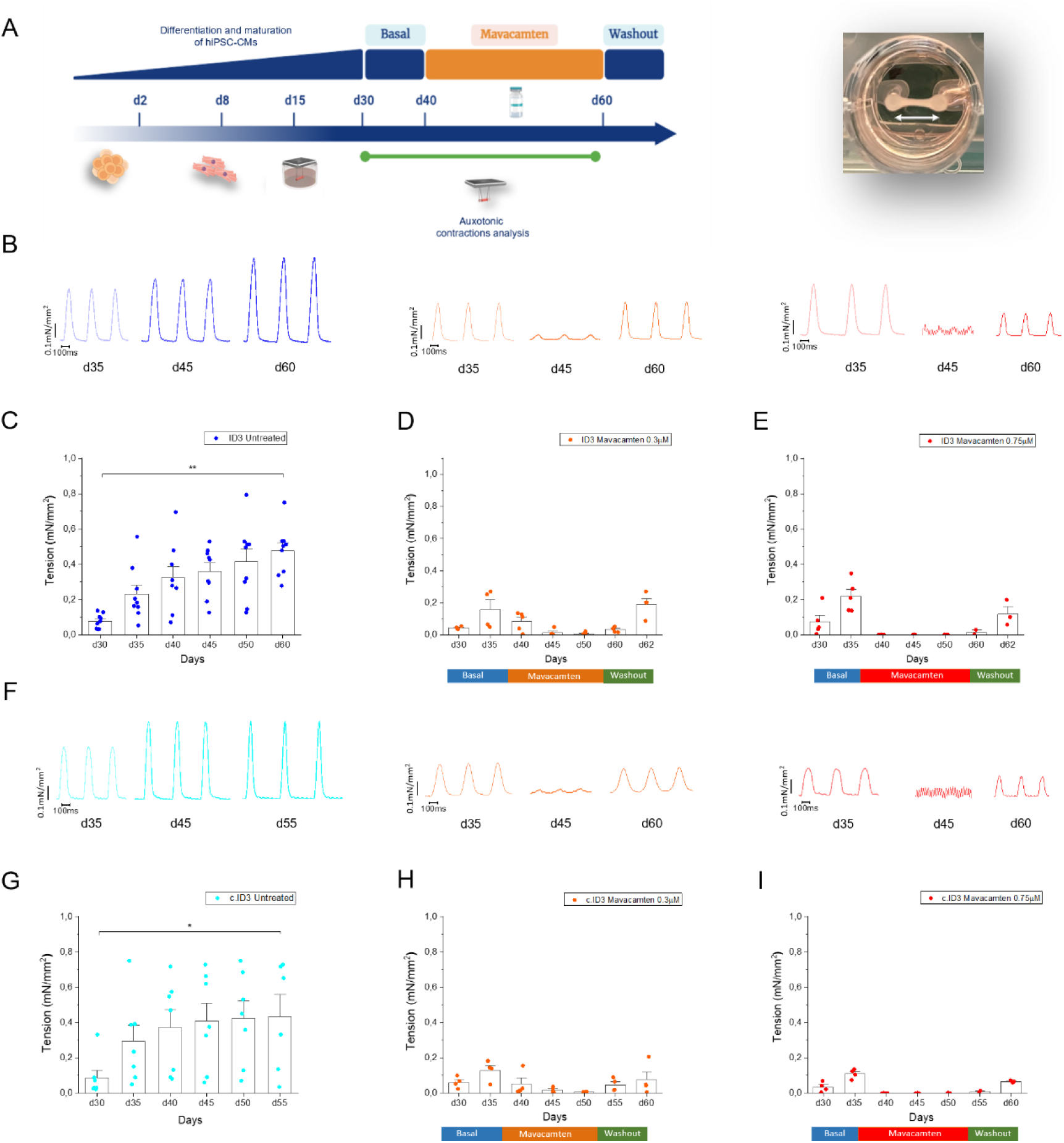
Spontaneous auxotonic contractions of c.ID3- and ID3-EHTs during chronic treatment with Mavacamten. **(A)** Schematic representation of chronic culture treatment of EHTs from day 30 to day 60 of differentiation. The scheme was created with BioRender.com. **(B)** Representative twitches of untreated and Mava-treated ID3-EHTs at d35, d45 and d60 p.d. **(C-D-E)** Spontaneous auxotonic tension of ID3-EHTs untreated (N=7; n=9;) and Mava-treated (Mava 0.3µM, N=6; n=4; Mava 0.75µM, N=5; n=5). **(F)** Representative twitches of untreated and Mava-treated c.ID3-EHTs at d35, d45 and d60 p.d. **(G-H-I)** Spontaneous auxotonic tension of c.ID3-EHTs untreated (N=5; n=7;) and Mava-treated (Mava 0.3µM, N=5; n=4; Mava 0.75µM, N=3; n=4) from day 30 to 60 of differentiation, measured at 37°C in RPMI/B27 culture medium (∼0.4mM of [ Ca^2+^]) at specific time points. Error bars ± SEMs. One-way analysis of variance (ANOVA) with a Tukey post-hoc test was used to compare the different time points. * p < 0.05 and ** p < 0.01 versus d30.

#### Effects of Mavacamten long-term treatment on the EHTs contractility under isometric fixed-end conditions

To confirm and extend these findings under more controlled and physiologically relevant conditions, we next evaluated EHT isometric contractile performance. After 20 days of treatment, the EHTs were detached from the pillars and mounted between a force transducer and a motor arm on a custom-built muscle mechanics apparatus. This setup allowed for the study of steady-state tension at imposed electrical stimulation frequencies. To monitor the time-course and consequence of drug removal, we compared the features of EHT contraction under isometric conditions just after removal of 0.3 µM Mavacamten (acute washout) and during the following 4-day recording. (**Figure 4A**). During the acute washout (d0), the tissues were perfused with the experimental solution (no Mavacamten) and during force recordings, contractions increased to a new steady state (after about 1h of perfusion). The ID3-EHT tension after chronic treatment with Mavacamten is reduced about 96% at day 0 post-Mavacamten (acute washout, 0.05±0.01 mN/mm²) and 63% at day 4 post-Mavacamten washout (0.46±0.14 mN/mm²) in comparison to the untreated tissues (1.24±0.2 mN/mm²) (**Figure 4B-C**). Furthermore, twitch duration was slightly accelerated in all treated groups ((d0) 293±10 ms, (d4) 277±21 ms) compared to the untreated groups (331±27 ms) (**Figure 4D**). Similar results are obtained at multiple pacing frequencies for both the c.ID3 **(Figure S1A-B**) and ID3 (**Figure S1C-D**). Although contractile tension remained lower in Mavacamten-treated EHTs compared to untreated controls, in ID3 tissues we observed an increase in contraction force by increasing the extracellular calcium concentration (from 0.5 mM to 4 mM), both at d0 (**Figure 4E-F**) and d4 post-washout (**Figure S2B- D**) as well as in c.ID3 (**Figure S2A-C**), suggesting a residual inotropic reserve, albeit not fully comparable to that of untreated groups. These results suggest a persistent effect of Mavacamten after long-term exposure. However, the recovery of force at day 0 and after washout appears slower than the rapid recovery observed in the acute experiment. Furthermore, to understand whether long-term treatment with Mavacamten had an effect at the electrophysiological level, we analysed the action potential duration (APD50, ms) in the same treated and untreated ID3-EHTs used for the previous experiment. Tissues were loaded with Fluovolt and then stimulated with regular pacing (1 Hz, 37°C in Tyrode’s solution). Compared to the untreated APD50, we observed no significant differences between treated (424±76 ms; 420±118 ms) and untreated EHTs (482±87 ms), indicating negligible impact on the electrophysiology following drug treatment at either concentration (**Figure S3A-B**).

**Figure 4.**
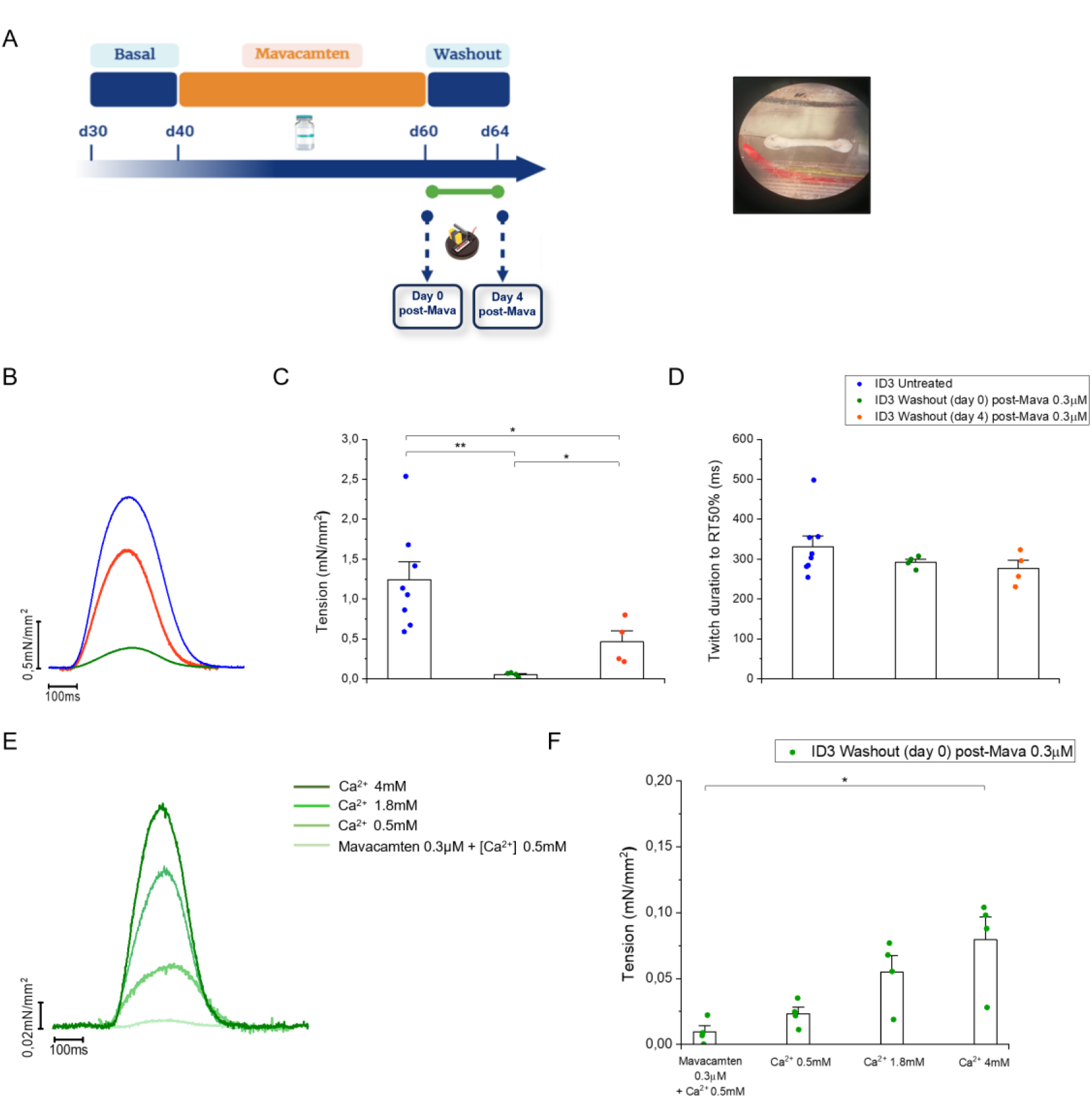
Contractile properties of EHTs after long-term Mavacamten exposure and washout. (A) Schematic representation of ID3-EHTs chronic treatment with Mavacamten 0.3 µM and washout at d0 and d4 after Mava-treatment. The scheme was created with BioRender.com. (B-C) Active tension and (D) twitch duration at 50% of relaxation of untreated ID3-EHTs (N=5; n=8) compared to EHT after treatment with Mavacamten (0.3 µM) at washout days 0 (N=3; n=3) and 4 (N=4; n=4). Recordings were performed at 1Hz constant pacing with 1.8 mM of [Ca^2+^]. (E-F) Active tension analysed at day 0 post-Mava washout by increasing the concentration of extracellular Ca^2+^ from 0.5 mM to 4 mM, under constant stimulation at 1 Hz. Error bars ± SEMs. One-way analysis of variance (ANOVA) with a Tukey post-hoc test was used. * p < 0.05 and ** p < 0.01.

### Sarcomere network evaluation in hiPSC-CMs

Given these results, suggesting disease-modification, we evaluated whether Mavacamten could have an effect on cardiac cell structure. Since both mutant and control EHTs showed similar functional results, we used a control hiPSC-CMs tagged with mEGFP alpha-actin for this series of experiments. Cardiomyocytes obtained after differentiation were seeded on nanopatterned surfaces to perform structural analysis on single cells. Similarly, to the EHT treatment protocol, clusters of alpha-actin- labelled cardiomyocytes were exposed to Mavacamten (0.3 µM and 0.75 µM) for 20 days. These were compared with untreated cells (DMSO) and with cells treated with 2,3-butanedione monoxime (BDM), another well-characterized, low-affinity and non-competitive inhibitor of myosin II, and with phenylephrine (PE), a selective alpha-1 adrenergic agonist that can induce cardiomyocyte hypertrophy (19) (**Figure S4A-E**). During cell culture, images of single CMs were acquired to assess the change in cell area at day 2 (**Figure S4F**), day 6 (**Figure S4G**), day 10 (**Figure S4H**) and day 15 (**Figure S4I**) post treatment. Finally, the sarcomere network was evaluated after the end of the treatment period and compared to untreated cells and after 4 days of washout (day 20 + 4 post treatment). From the analysis performed at various days of maturation, the drug-treated cardiomyocytes showed a reduction in cell area during long-term exposure to Mavacamten. Similarly, the reduction in cell area was observed with BDM although cardiomyocytes did not survive long after days of culture, differently to Mavacamten. In contrast, PE-treated CMs did not show significant changes in the cell area over time but remained larger compared to the Mavacamten- and BDM- treated groups. Furthermore, we observed the same results by monitoring the area of the mutated cardiomyocytes (ID3) treated with both Mavacamten and Aficamten (**Figure S5A**), a second- generation cardiac myosin inhibitor (18–20), using the same concentrations (0.3 and 0.75 µM) at day 2 (**Figure S5B**), day 8 (**Figure S5C**), day 15 (**Figure S5D**) post treatment. At the end of chronic treatment (day 50 post differentiation), confocal imaging was used to quantify the percentage of sarcomere filaments per cell area in both treated and untreated GFP-labelled cardiomyocytes (**Figure 5A-E**). Quantitative analysis revealed a reduction in sarcomere content in Mavacamten-treated cells to 0.3 µM (29.82±0.50 %) and 0.75 µM (25.98±0.59 %) compared with untreated CMs (37.04±0.48 %). We also observed that after 4 days of washout, sarcomere density was slightly increased (Mava 0.3 µM 32.08±0.66 %; Mava 0.75 µM 32.30±0.61 %) compared with cells in which the drug was not removed (**Figure 5F**). The incomplete restoration of myofibril density after 4-day washout may explain the low force observed after washout in EHTs treated chronically with Mavacamten. In addition, the mean cell area was also significantly reduced in the Mavacamten-treated α-actinin labelled CMs (Mava 0.3 μM, 1971±124 μm^2^; Mava 0.75 μM, 1787±142 μm^2^) compared with the untreated cell group (2819±206 μm^2^), with a slight increase after washout (**Figure 5G**). Finally, to understand whether the reduction in contractility caused by Mavacamten may influence the normal turnover of myofibril formation and disassembly during cell maturation, we treated single CMs with both Mavacamten concentrations at an advanced stage of cell maturation (day 55 post differentiation).

**Figure 5.**
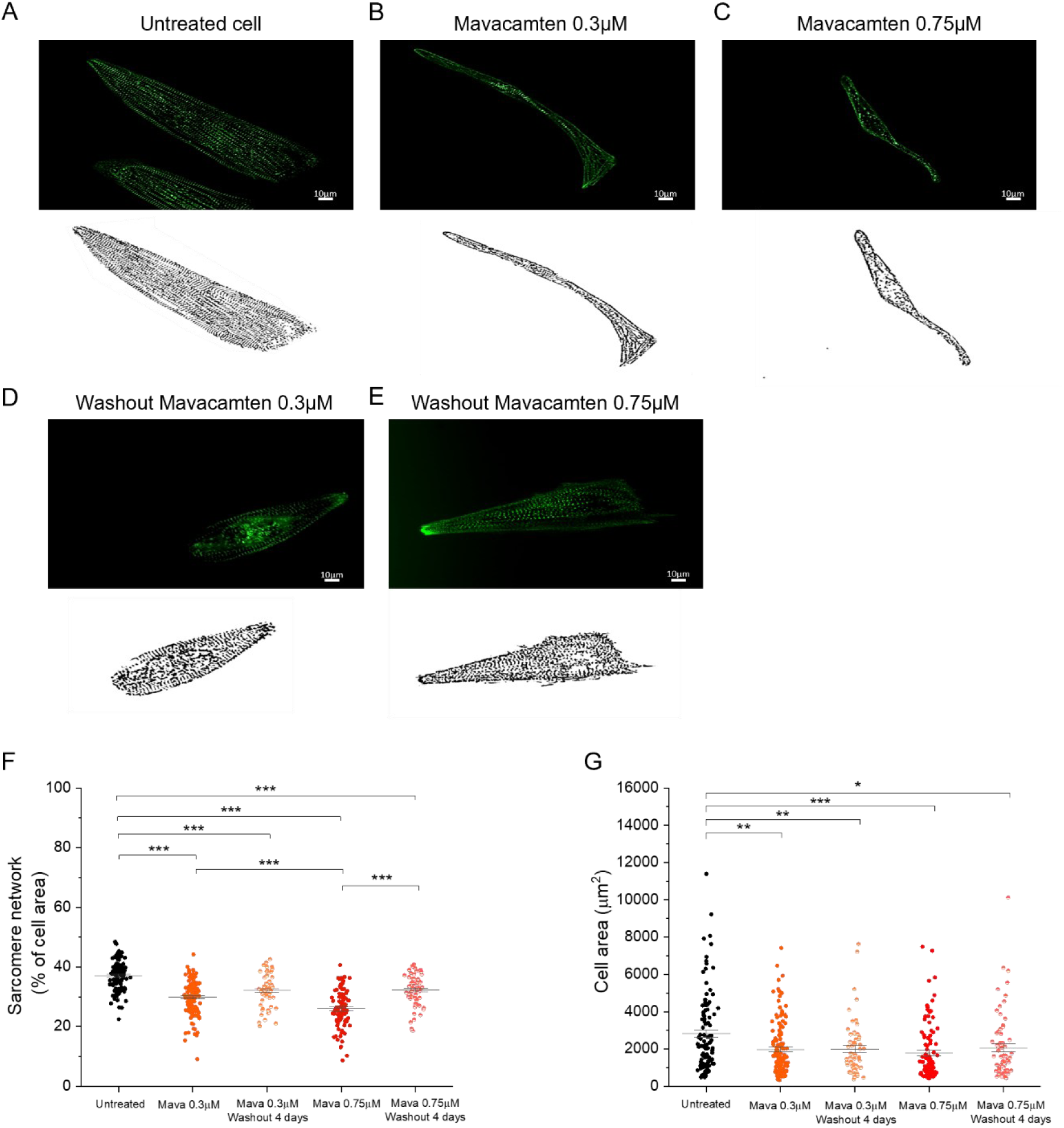
Sarcomere density evaluation in cardiomyocytes derived from mEGFP-tagged α- actinin hiPSC treated until day 50 of maturation. (A-E) Representative confocal images recorded at 60X magnification under different treatment conditions and binary images of the α-actinin network showing the difference in sarcomere density in treated versus untreated cell groups. (F) Density of the sarcomere network and (G) cell area analysis in labelled hiPSC-CMs at day 50 of maturation. Untreated cells (N=2; n=105) were compared with CMs treated with Mavacamten at 0.3 µM (N=2; n=133) and 0.75 µM (N=2; n=106) for 20 days and treated cells subjected to a 4-day washout (N=2; n=62; n=68). The mean sarcomere density is significantly reduced in Mava-treated CMs (29.82 % ± 0.50; 25.98 % ± 0.59) compared to untreated CMs (37.04 % ± 0.48). Error bars ± SEMs. Statistical analysis was performed using one-way ANOVA, with a Tukey post-hoc test with statistical significance set at * p < 0.05, ** p < 0.01, *** p<0.0001.

After only 24 hours of drug exposure (**Figure 6A-C**), confocal imaging again showed a reduction in sarcomere network density (**Figure 6D-E**) and cell area (**Figure 6F-G**), suggesting that Mavacamten exerts its effects through myofibril disassembly. Consistently, EHTs at an advanced stage of maturation (day 55 post differentiation) treated for 24h with Mavacamten 0.3 µM showed persistent reduction in active tension compared to untreated group (0.32±0.10 mN/mm² vs. 1.24±0.22 mN/mm²) (**Figure S6A-C**). Overall, these results indicate that the sustained reduction in active tension observed in Mavacamten-treated EHTs results from an increased rate of myofibril disassembly relative to formation, rather than interference with EHT maturation.

**Figure 6.**
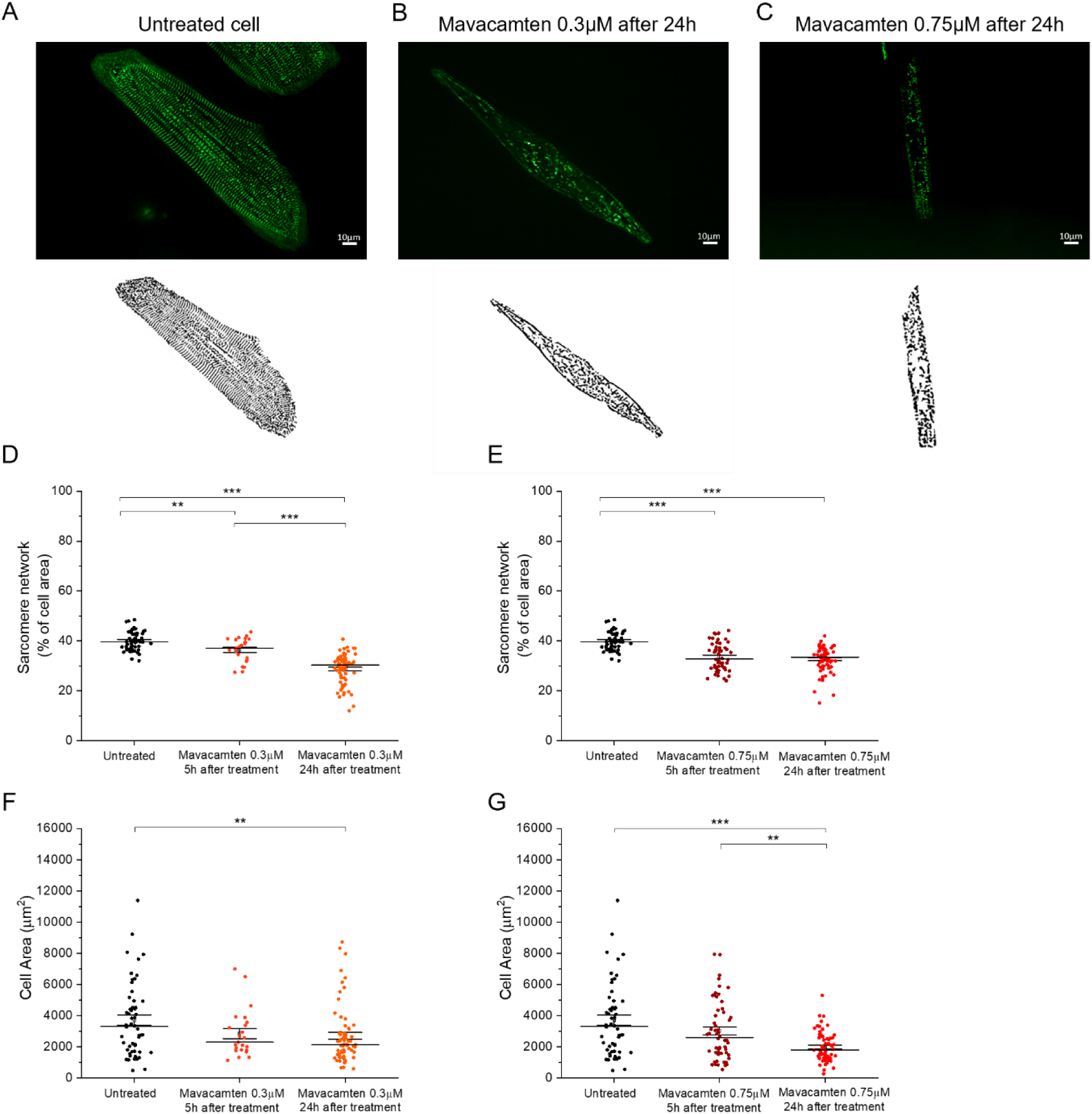
Sarcomere density evaluation in mature α-actin-labelled cardiomyocytes (d50 p.d.) treated with Mavacamten for 5h and 24h. (A-B-C) Fluorescence and binary images of alpha-actin-labelled cardiomyocytes (d50 p.d.) untreated and treated with Mavacamten for 24 h. (D-E) Sarcomere network density and (F-G) cell area analysis in untreated hiPSC-CMs (N=2; n=54) compared to treated CMs with Mavacamten at 0.3 µM (N=2; n=23; n=68) and 0.75 µM (N=2; n=56; n=65) for 5 h and 24 h. The mean sarcomere density is significantly reduced in Mava-treated CMs after 24 h (28.75 % ± 0.73; 32.64 % ± 0.64) compared to untreated CMs (40.05 % ± 0.51). Error bars ± SEMs. Statistical analysis was performed using one-way ANOVA, with a Tukey post-hoc test with statistical significance set at * p < 0.05, ** p < 0.01, *** p<0.0001.

### Transcriptome analysis in the EHTs

To investigate the effects of Mavacamten at the molecular level we performed a transcriptome analysis on EHTs at day 60 from ID3 e c.ID3 cell lines. Expression differences were assessed between isogenic untreated vs. mutated untreated and mutated untreated vs. mutated treated with Mavacamten 0.3 µM and 0.75 µM. This analysis identified 487 differentially expressed genes (353 up, 134 down) for the mutation versus isogenic control cell lines, predominantly linked to immune response signatures (**Figure 7A**), likely reflecting heightened cellular stress and fibroblast presence. We also observed enrichment for cardiac muscle tissue development (GO:0048738), calcium ion homeostasis (GO:0006874), and extracellular matrix organization (GO:0030198; **Figure 7B**). A Gene-Concept Network highlighted global upregulation of these pathways (**Figure 7D**). To confirm expected disease-relevant transcriptional signatures, we specifically examined the expression of key HCM markers, including MYBPC3, MYL2, TNNI3; calcium-handling genes such as CASQ2 and ATP2A2; fibrosis-associated genes (LOX, LOXL2, COL5A3, COL12A1, TNXB, DPT); and signaling factors (STC1, TBX20, PROX1), and visualized them in a heatmap (**Figure 7C**). Supporting the pathophysiological relevance of the model, we observed reduced expression of ATP2A2 (SERCA2) and repolarizing potassium channel genes (KCNIP2, KCNJ2) in ID3 EHTs, mirroring changes previously reported in human HCM samples (20). In line with previous observations, in the ID3 cell lines and the same patient derived tissue (14,15), downregulation of the MYBPC3 gene was observed in our ID3-EHTs, supporting the c-MyBPC haploinsufficiency hypothesis (**Figure 7C**). Furthermore, Gene Set Enrichment Analysis (GSEA), performed using the Hallmark gene sets from the Molecular Signatures Database (MSigDB), showed an upregulation of oxidative phosphorylation genes in mutant samples (ID3) compared to the isogenic control (c.ID3) (**Figure S7**). Comparing untreated ID3 with the 0.3 μM and 0.75 μM treatments, differential expression analysis revealed 2461 (1018 upregulated, 1443 downregulated) and 1453 (558 upregulated, 895 downregulated) significantly altered genes, respectively. To disentangle concentration-dependent from concentration-independent effects, we conducted overrepresentation analysis on each gene set and cross-referenced the results to define “common” and “exclusive” activities (**Figure 8**) (**Table 1**). The common activities included ATP synthesis (GO:0006754, GO:0046034, GO:0015986), extracellular matrix organization (GO:0030198, GO:0043062), and sarcomere development (GO:0045214), supporting a dual role for Mavacamten in modulating sarcomere function and exerting metabolic and anti-fibrotic benefits (**Figure S9**). In contrast, most cardiac-specific processes fell into the exclusive category, suggesting that higher concentrations drive more targeted effects on cardiomyocyte biology (**Figure S8**). Gene set enrichment analysis (GSEA) against KEGG pathways corroborated these findings: both treatments showed perturbation of the cardiac muscle contraction pathway (hsa04260; **Figure S11– S12**). Exclusive to the 0.3 μM condition were endoplasmic reticulum processing (hsa04141) (**Figure S13**) suggesting increase degradation activities of protein synthesis, whereas the 0.75 μM showed upregulated pathways involved in protein turnover and mechanotransduction (SGCE, SGCD, TNNI3, TNNT2, ITGA4, TPM2) (**Table 1**, **Fig. S9**), in the KEGG hypertrophic cardiomyopathy pathway (hsa05410) (**Figure S10**). Notably, in the 0.75 μM cluster, the cardiac muscle contraction pathway (hsa04260) showed, however, that myosin is downregulated (**Figure S11)**. In addition, PI3K-Akt signaling (hsa04151) genes implicated in inflammation and fibrosis (e.g., COL1A1, COL6A2, COL4A1, FGF18) were downregulated (**Table 1**). Together, these transcriptional changes provide a molecular framework linking long-term effects of Mavacamten to structural remodelling of sarcomere and cytoskeleton, improved contractile function and attenuation of fibrotic and inflammatory signaling in ID3 EHTs.

**Figure 7.**
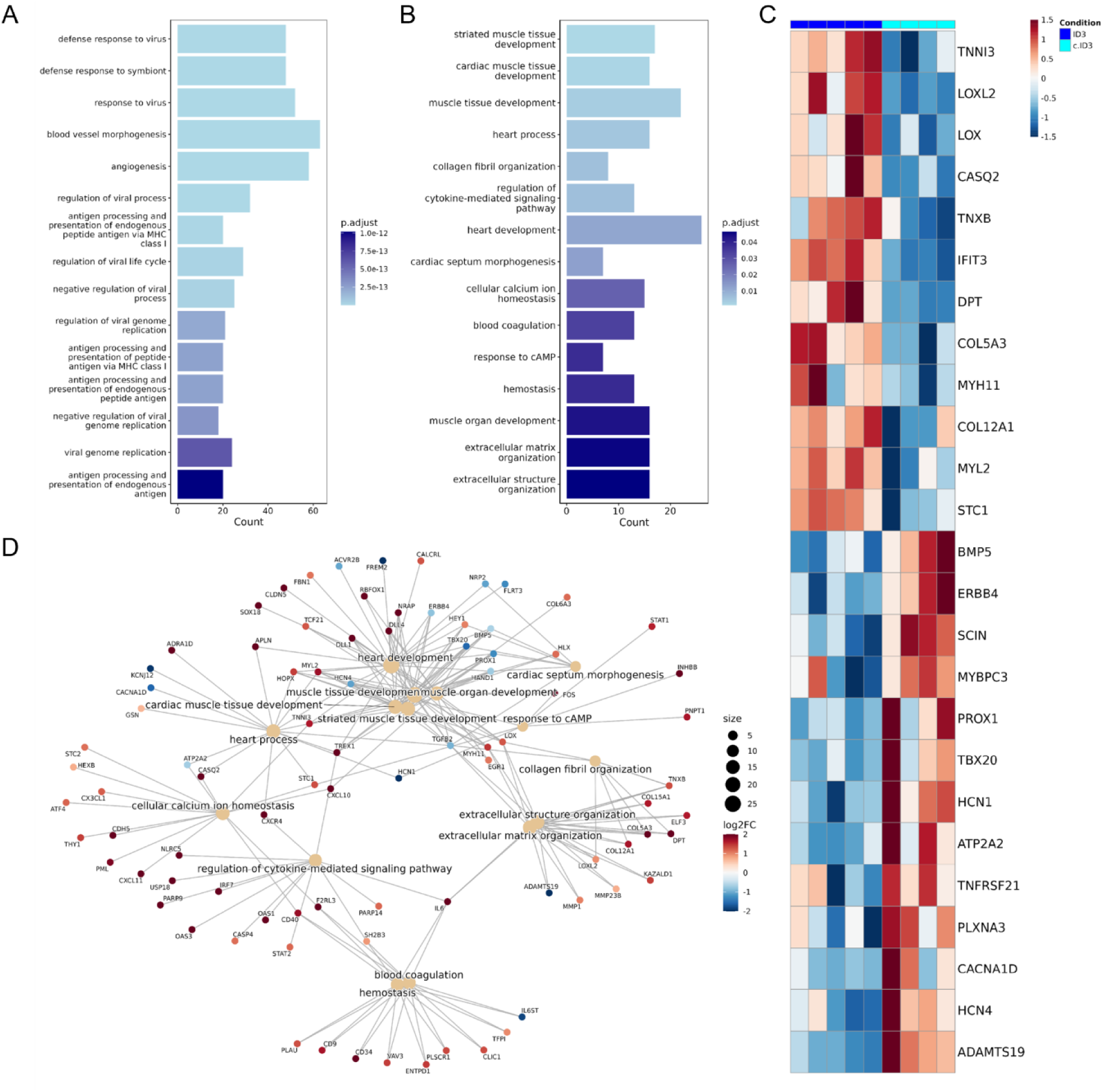
Transcriptomic analysis performed on c.ID3- and ID3-EHTs. (A) Gene Ontology (GO) enrichment analysis of differentially expressed genes (DEGs) between untreated HCM and isogenic samples (ID3 N=5, n=5; c.ID3 N=4, n=4). The bar plot displays the top 15 significantly enriched GO terms ranked by adjusted p-value (padj); (B) Subset of GO terms from panel A, filtered to emphasize pathways of particular interest. (C) Heatmap of a subset of DEGs (padj < 0.05) selected based on known markers of interest. The color scale reflects the gene expression based on the log2 fold change (LFC), red=upregulation and blue=downregulation; (D) Gene ontology (GO) enrichment network of DEGs. GO terms (yellow nodes), previously identified in the bar plot in panel B, are connected to their associated DEGs (colored nodes), filtered for significance (adjusted p-value < 0.05). Node color indicates log2 fold change (log2FC) in gene expression, while node size reflects the number of genes linked to each GO term.

**Figure 8.**
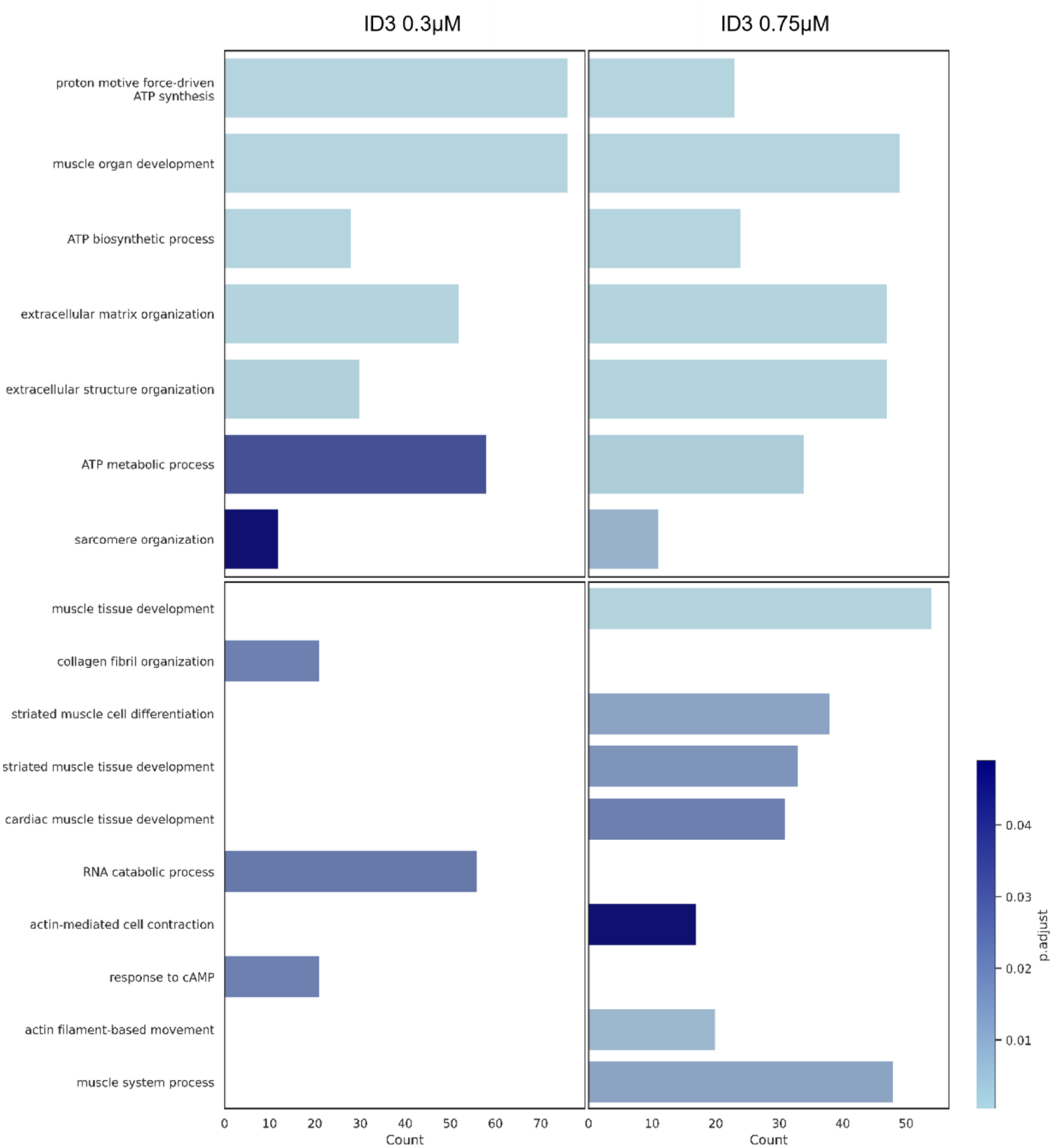
Enrichment levels of common (top panels) and exclusive (bottom panels) biological processes in ID3 EHTs treated with Mavacamten at 0.3 µM and 0.75 µM. Bar plot showing processes represented by gene counts (x-axis), and color intensity adjusted p-values (P.adjust), as shown in the legend; (0.3 µM n=3; 0.75 µM n=3)

**Table 1.**
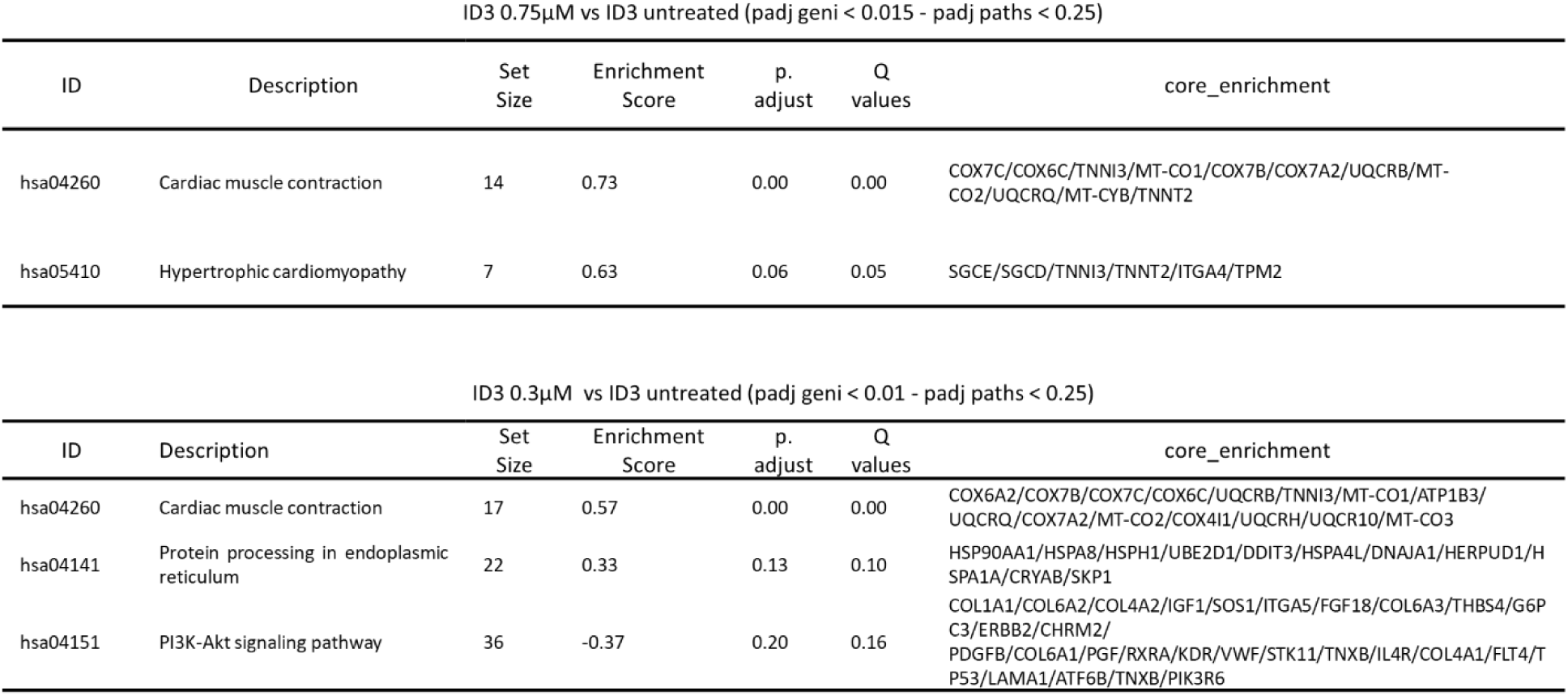
KEGG pathway enrichment analysis. Results for two comparisons between untreated mutant cells (n=4) and those treated with Mavacamten at two different concentrations (0.3 µM and 0.75 µM) (n=3). Each row represents a significantly enriched KEGG pathway, with bars indicating statistical significance (expressed as -log10(p-value)) for each comparison.

| ID3 0.75µM vs ID3 untreated (padj geni < 0.015 - padj paths < 0.25) |  |  |  |  |  |  |
| --- | --- | --- | --- | --- | --- | --- |
| ID | Description | Set Size | Enrichment Score | p. adjust | Q values | core_enrichment |
| hsa04260 | Cardiac muscle contraction | 14 | 0.73 | 0.00 | 0.00 | COX7C/COX6C/TNNI3/MT-CO1/COX7B/COX7A2/UQCRB/MT-CO2/UQCRQ/MT-CYB/TNNT2 |
| hsa05410 | Hypertrophic cardiomyopathy | 7 | 0.63 | 0.06 | 0.05 | SGCE/SGCD/TNNI3/TNNT2/ITGA4/TPM2 |

| ID3 0.3µM vs ID3 untreated (padj geni < 0.01 - padj paths < 0.25) |  |  |  |  |  |  |
| --- | --- | --- | --- | --- | --- | --- |
| ID | Description | Set Size | Enrichment Score | p. adjust | Q values | core_enrichment |
| hsa04260 | Cardiac muscle contraction | 17 | 0.57 | 0.00 | 0.00 | COX6A2/COX7B/COX7C/COX6C/UQCRB/TNNI3/MT-CO1/ATP1B3/UQCRQ/COX7A2/MT-CO2/COX4I1/UQCRH/UQCR10/MT-CO3 |
| hsa04141 | Protein processing in endoplasmic reticulum | 22 | 0.33 | 0.13 | 0.10 | HSP90AA1/HSPA8/HSPH1/UBE2D1/DDIT3/HSPA4L/DNAJA1/HERPUD1/HSPA1A/CRYAB/SKP1 |
| hsa04151 | PI3K-Akt signaling pathway | 36 | -0.37 | 0.20 | 0.16 | COL1A1/COL6A2/COL4A2/IGF1/SOS1/ITGA5/FGF18/COL6A3/THBS4/G6PC3/ERBB2/CHRM2/PDGFB/COL6A1/PGF/RXRA/KDR/VWF/STK11/TNXB/IL4R/COL4A1/FLT4/TP53/LAMA1/ATF6B/TNXB/PIK3R6 |

## Methods

An expanded Methods section is available in the *Supplemental Material*.

### HiPSC generation and cardiac differentiation

PBMCs from a male HCM patient harboring the MYH7 c.772G>A mutation were reprogrammed using the CytoTune®-iPS 2.0 Sendai system (14,15). Clones were selected via flow cytometry and verified by karyotyping and Sanger sequencing (14,15). A homozygous isogenic control line (c.ID3) was created using CRISPR/Cas9 editing. hiPSCs were differentiated into cardiomyocytes using a monolayer protocol and Cardiomyocyte Differentiation Kit. For imaging analyses, mEGFP-tagged α-actinin-2 hiPSCs were cultured on nanopatterned surfaces.

### Engineered heart tissue (EHT) generation and force measurements

EHTs were generated using fibrin scaffolds containing 1×10⁶ hiPSC-CMs and cultured on PDMS racks. Spontaneous and isometric contractile force was measured optically (post deflection) and with force transducers under variable calcium concentrations (0.5–4 mM), pacing rates (0.2–2.5 Hz), and drug conditions. Mavacamten was applied chronically (0.3–0.75 μM) for 20 days, followed by washout in select experiments.

### Imaging and electrophysiology

Sarcomere density and cell area were quantified by ImageJ using ridge detection. Action potentials (APs) and calcium transients (CaTs) were recorded from day 60 EHTs using Fluovolt and Cal630 dyes. This was done under Tyrode perfusion at 37 °C, with blebbistatin or mavacamten, and imaging was done using confocal microscopy during field stimulation (0.1–2 Hz).

### Transcriptomics

RNA was extracted from EHTs (treated and untreated) using QIAzol and RNeasy kits. RNA-seq was performed by Breda Genetics and analyzed using Salmon, DESeq2, and batch-correction tools (ComBat, RUVSeq). DEGs were defined as adj. p < 0.05 and |log₂FC| > 0.5. Gene set enrichment analyses (GSEA) were performed using clusterProfiler against GO and KEGG databases.

### Statistics

Data are presented as mean ± SEM. Group comparisons were made using a two-tailed Student’s *t* test and one-way ANOVA with Tukey post-hoc test. Significance was defined as p < 0.05. Analyses were conducted in OriginPro.

## Discussion

This study provides mechanistic insight into the persistent effects of long-term Mavacamten exposure on sarcomere organization, cardiomyocyte transcriptomics, and contractile function in an in vitro human hypertrophic cardiomyopathy (HCM) model. Follow-up studies in the EXPLORER HCM trial have shown evidence for the first time that a compound designed for HCM can cause cardiac remodeling with reduced left ventricular (LV) mass in a short period of time (30 weeks) (11,12), which is consistent with the pre-clinical disease-modifying effects observed in transgenic mice (3). While initial hypotheses attributed this remodeling primarily to relief of afterload via LVOT gradient reduction (12), our findings suggest that Mavacamten may also exert direct structural effects on sarcomere architecture independent of the cardiac genetic background. The *MYBPC3*:c.772G>A hiPSC line exhibits a distinctive hypertrophic cardiomyopathy (HCM) phenotype, characterized by an apparent preserved twitch. This phenotype is the result of a compensatory mechanism involving altered sarcomere energetics, and makes the line a suitable model for investigating long-term Mavacamten. Using engineered heart tissues (EHTs) derived from the patient hiPSC line and its isogenic control, we show that chronic exposure to Mavacamten leads to dose and time dependent reductions in sarcomere density and force production, without affecting the electrophysiological profile. Transcriptomic analyses suggests additional antifibrotic and metabolic effects, suggesting that Mavacamten benefits extend beyond sarcomere energetics. These findings provide a deeper understanding of how sustained myosin inhibition may contribute to reverse remodeling in HCM and raise important considerations for timing, dosing, and patient selection.

### Mavacamten induces dose- and time-dependent structural remodeling

Our study demonstrates that chronic treatment with Mavacamten induces a progressive and dose dependent reduction in cardiomyocyte size and sarcomere density. These structural changes underlie the observed decrease in twitch force and sarcomeric energetics in both *MYBPC3*-mutant (ID3) and isogenic control (c.ID3) engineered heart tissues (EHTs). Chronic treatment produced partial (0.3 μM) or near-complete (0.75 μM) reductions in twitch force, and led to recovery after a 4-day washout period.

Furthermore, long-term Mavacamten treated EHTs preserved a positive inotropic response upon addition of extracellular calcium, suggesting that calcium handling is not affected (inotropic reserve) and myosin heads sequestered in the off state(s) induced by Mavacamten are recruitable after increase of calcium binding to thin filament in agreement with previous findings under β-adrenergic stimulation and length-dependent activation (9,17,21). Transcriptomic data showed no major changes in thick or thin filament gene expression, suggesting that sarcomeric remodeling may occur through post-transcriptional and/or removal of mechanical feedback mechanisms consecutive to force reduction. Similar results on cell structure were obtained with other myosin inhibitors, including blebbistatin (22), 2,3-butanedione monoxime (BDM), and Aficamten (**Figure S4 and S5**), a second- generation cardiac myosin inhibitor (23–25). One potential mechanism may therefore be related to the reduction of force (hypocontractility) leading to reduced organization of sarcomere and cytoskeletal proteins rather than a specific effect of Mavacamten. The reduction in force observed with both Mavacamten concentrations over time demonstrates the drug’s dose-dependent effect without causing noticeable cell death. This is different from what we observed following prolonged exposure of cells to BDM (**Fig. S4**), which further reinforces the safety of Mavacamten. Notably, myofibril density following sustained force depression (0.75 μM) was restored to the same extent as partial force depression (0.3 μM) after washout.

### Mechanistic insights into sarcomere regulation

Biophysical data support a dual-phase model of Mavacamten action: an immediate (short-term) reduction in active force through the removal of myosin from the pool that is available to participate in contractile force generation (6), followed by a slower (long-term), structural remodeling of the sarcomere network. Indeed, when EHTs from both *MYBPC3*:c.772G>A variant and its isogenic corrected counterpart were treated for a short period of time (minutes), force decreased, and a new steady state was reached. This could be completely restored after washout. Conversely, EHTs chronically exposed to Mavacamten exhibited slower force recovery during a four-day washout period. In untreated conditions, sarcomere network was estimated to be 40% of the cell area in single hiPSC-CMs, in agreement with previous findings (26). We observed that sarcomere density decreased to approximately 30 and 25 % of cell area (with 0.3 and 0.75 μM respectively) in long-term treated hiPSC-CMs that had undergone long-term treatment. This finding is consistent with the residual reduction of active force after four days of washout. In parallel, transcriptomic data revealed reduced expression of pathways involved in protein folding and upregulation of endoplasmic reticulum (ER)- associated degradation (ERAD) proteins, which targets misfolded proteins of the endoplasmic reticulum for ubiquitination and subsequent degradation of proteasome protein. These pathways may contribute to the enhanced clearance of myosin and other sarcomere proteins. In accordance with sarcomere structure disassembly, hiPSC-CMs on nanopatterned surfaces after reaching the highest level of structural maturity (day 50 p.d.) demonstrated reduced myofibril density and force within 24 hours. In adult sarcomeres, protein turnover between assembly and disassembly is nearly balanced (27), whereas hiPSC-derived cardiomyocytes, even when matured to adult-like sarcomere density, remain developmentally biased toward assembly. The observation that Mavacamten reduces sarcomere density therefore reinforces the notion that its effect favors rapid disassembly rather than slowed assembly that may impact maturation, further supporting the value of this model for mechanistic studies of sarcomeric remodeling. For instance, it was proposed that Mavacamten directly enhances the mobility of myosin out of the thick filament increasing its solubility, and a stabilization of a compact/folded conformation of the molecules (28). This mechanism is compatible with a higher rate of protein degradation, which may contribute to a long-term net reduction in sarcomere content observed over time. Finally, since the gene expression program of major thick- and thin-filament proteins is unaltered, this may facilitate the rapid reassembly of myofibrils after washout.

### Sarcomere remodelling without disruption of Excitation–Contraction Coupling

Despite marked structural changes, key aspects of Excitation-Contraction Coupling (ECC) were preserved. Action potential duration (APD) was unaffected across treated groups, indicating that the electrophysiological profile remains altered in the HCM cell line. On the other hand, twitch duration showed only a trend towards acceleration, a variation that was more evident in other studies using similar models (17,29). This suggests that the observed reduction of twitch duration is primarily mechanical rather than electrical in nature. Variability exists, which may reflect differences in tissue homogeneity. The mild shortening in twitch kinetics observed may be attributed to faster cross-bridge detachment (8), thereby contributing to the shortened relaxation phase. In this context, it is also plausible that the reduction in sarcomere network and the decreased number of available myosin heads may contribute to the slightly faster twitch kinetics, reflecting reduced sarcomere resistance to relaxation.

These findings align with clinical observations of preserved or improved diastolic function following Mavacamten treatment (3,6) and supports the idea that Mavacamten optimizes sarcomere performance by tuning the balance between force generation and relaxation without altering E-C coupling dynamics at the cell level. Moreover, while APD remained stable in our in vitro model, previous clinical studies have reported reduced QT dispersion after long-term therapy (30). This discrepancy can be explained by the fact that QT dispersion reflects repolarization heterogeneity at the organ level, not absolute APD.

### Reduction of pro-fibrotic and upregulation of metabolic pathways

In addition to all these findings, transcriptomic analysis of HCM-EHTs also revealed the presence of a chronic inflammatory profile that can trigger activation of fibrotic processes, including the upregulation of COL5A3, COL32A2, and LOXL2. This response is further stimulated by overexpression of IL-6, a key molecular mediator of left ventricular hypertrophy and myocardial fibrosis (31). These findings are consistent with previous single-cell comparative transcriptomic studies in both human and animal HCM samples (3,32) likely due to the presence of cardiac fibroblasts in the EHTs. Transcriptomic analysis revealed that Mavacamten downregulates fibrosis- associated gene pathways, including those involved in TGF-β signaling and extracellular matrix organization. This potential antifibrotic activity may contribute to improved diastolic function in the patient myocardium and a less arrhythmogenic myocardial substrate, potentially explaining clinical findings of reduced fibrosis, reduced QT dispersion and improved electrical homogeneity. Concurrently, HCM-EHTs showed increased oxidative phosphorylation pathways, which is consistent with altered energy metabolism in HCM (33). Altered cell metabolomics is another hallmark, particularly, of genotype-positive HCM patients (34). With Mavacamten treatment, metabolic profiling showed enhanced mitochondrial efficiency such as ATP metabolic and biosynthetic processes. These changes suggest that Mavacamten not only reduces mechanical strain but also improves cellular resilience and energy balance, supporting its potential in restoring cellular energetics in HCM. These mechanisms support the hypothesis that Mavacamten alleviates sarcomere- driven bioenergetic imbalance and can be particularly efficient in genotype-positive HCM (35–37). In this regard, we previously observed that upon exposure to Mavacamten, the HCM (ID3) hiPSC- CMs, but not the isogenic c.ID3 line, reduced the basal respiration and ATP production of the patient hiPSC-CMs, again suggesting greater benefit in reducing the ATPase activities (15).

### Translational implications

Our findings provide a novel mechanistic explanation for the reduction in left ventricular (LV) wall thickness and mass index observed in clinical trials after 18–30 weeks of Mavacamten treatment (11,12). Rather than reflecting simple afterload reduction, these changes seem to reflect structural remodeling of the sarcomere network.

Importantly, washout experiments demonstrate that this remodeling is reversible, with contractile force significantly recovering within days after drug removal. This mechanistic model aligns with clinical evidence demonstrating a positive impact on obstructive HCM (oHCM), reducing myocardium hypercontractility and wall thickness ameliorating LV outflow tract (LVOT) obstruction. The structural and functional changes induced by Mavacamten are dynamic and adaptable, supporting the concept of therapeutic reverse remodeling in oHCM. These findings also align with clinical experience in obstructive HCM, where reductions in left ventricular ejection fraction (LVEF) below 50% are uncommon. Our data indicate reverse remodeling, even in cases of excessive contractile suppression, upon washout, underscoring the favorable safety profile of Mavacamten. This further strengthens the translational value of our in vitro model, which provides mechanistic insight consistent with patient outcomes. However, it is important to note that our findings are derived from an in vitro model using sustained Mavacamten exposure, designed to probe the mechanistic effects. While informative, the dose and timing may not directly mirror the pharmacokinetics or tissue exposure experienced by patients with oHCM receiving long-term Mavacamten therapy.

Notably, despite the excellent safety profile observed in patients up to 180 week in the Explorer Long term extension (38), the potential for maladaptive structural remodeling suggested by our findings should be considered, in pre-hypertrophic or very early HCM. Properly designed studies are needed to assess the long-term effects of myosin heavy chain inhibition in early-stage HCM; however, these appear challenging due to the sample size and observation periods required. In this context, patient- specific hiPSC-CM models provide a powerful predictive platform to explore long-term safety and efficacy, bridging mechanistic insights with translational relevance.

## Author contributions

All experiments were conducted at the University of Florence. M.L. and J.M.P. designed the overall research study and wrote the manuscript. M.L., L.G., and J.M.P. conducted electro-mechanical experiments, acquired fluorescence and confocal imaging data, and analyzed the overall data. J.M.P. provided material and reagents. S.S. and M.R. provided hiPSC lines. R.S. and E.M. acquired and analyzed transcriptomic data. R.S. and A.M. assisted with bioinformatics training. B.S. managed the database. I.D.C. conducted mechanical experiments. M.L. and V.S. extracted RNA for sequencing.

V.S. and E.C. assisted with RNA extraction training. L.S., E.C., M.R., I.O., C.T., C.P., R.C., C.F. and J.M.P. provided guidance, support, and expertise to researchers. All authors critically reviewed the manuscript and approved the final version for publication.

## Acknowledgments

This work was supported by grants to Horizon Europe HORIZON-HLTH-2023-TOOL-05 SMASH-HCM grant number 101137115 (J.M. Pioner, C. Poggesi, R. Coppini, C. Ferrantini and I. Olivotto) and European Union’s Horizon 2020 under grant agreement No. 777204 SILICOFCM (M. Regnier and C. Poggesi). M. Langione is currently supported by a postdoctoral fellowship funded by the Ministero dell’Università e della Ricerca (MUR) under the Progetti di Rilevante Interesse Nazionale (PRIN) 2022 PNRR program, titled ’MALADAPT’ under the grant number 20223L2C9N (J.M. Pioner)

---

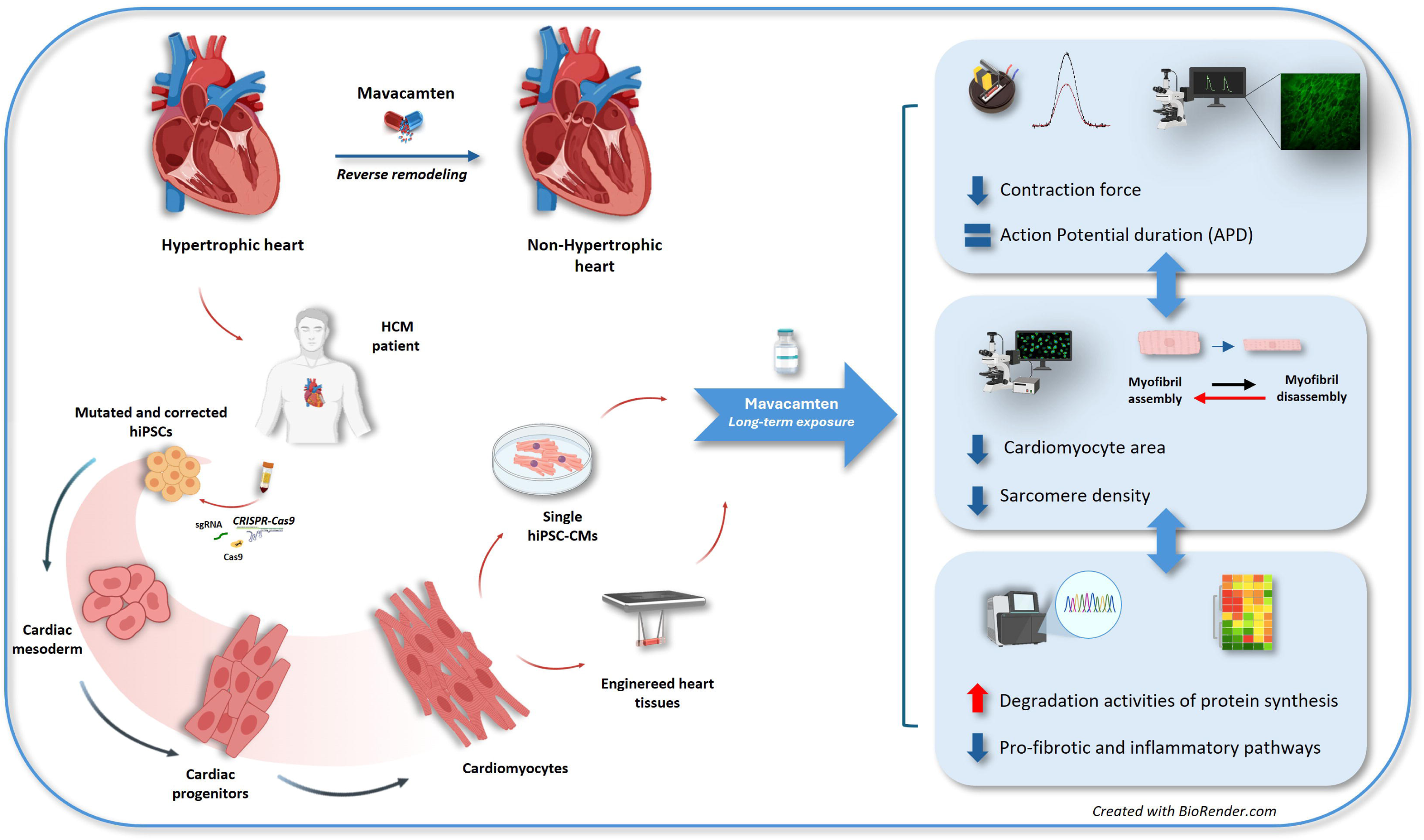

