## Supplemental material for "Long-term mavacamten exposure reduces force and sarcomere density in a hiPSC model of hypertrophic cardiomyopathy"

**Conflict of interest:** Iacopo Olivotto has received advisory board fees/research grants from Bristol Myers Squibb, Cytokinetics, Sanofi Chiesi, Genzyme, Amicus, Bayer, Tenaya, Rocket Pharma and Edgewise and Lexeo. Sonette Steczina was a former graduate student at the University of Washington at the time of this work and is currently an employee at Cytokinetics

**Table of contents:**

Supplementary materials and methods ……………………………………………...page S3
Figure S1……………………………………………………………………….....page S6
Figure S2……………………………………………………………………….....page S7
Figure S3……………………………………………………………………….....page S8
Figure S4……………………………………………………………………….....page S9
Figure S5……………………………………………………………………….....page S10
Figure S6……………………………………………………………………….....page S11
Figure S7……………………………………………………………………….....page S12
Figure S8……………………………………………………………………….....page S13
Figure S9……………………………………………………………………….....page S14
Figure S10……………………………………………………………………….page S15
Figure S11……………………………………………………………………….page S16
Figure S12……………………………………………………………………….page S17
Figure S13……………………………………………………………………….page S18
References……………………………………………………………………….page S19

### EXPANDED MATERIALS AND METHODS

##### *Stem cell line generation*

The protocols for stem cell line generation from human peripheral blood mononuclear cells (PBMCs), reprogramming into iPSCs, and iPSC CRISPR/Cas9 gene editing have been recently published (1,2). Briefly, PBMCs were isolated using a standard isolation method from patient blood samples collected at Careggi Hospital (Florence, Italy). PBMCs were isolated by density gradient centrifugation Ficoll (sodium diatrizoate, polysaccharides, and water, density of 1.08 g/mL) in equal volume of whole blood diluted in PBS and centrifuged for 40 minutes at 400-500 x g without brake. The PBMCs were collected at 2 x $10^6$ cell/ml in 20% dimethyl sulfoxide (DMSO) and 90% Fetal Bovine Serum (FBS) (Life Technologies). The cell line (PBMCs) obtained from a patient carrying the c.772G>A mutation was reprogrammed into iPSCs utilizing the CytoTune®-iPS 2.0 Sendai Reprogramming Kit (Gibco). A representative patient line (ID3) was genome edited using CRISPR/Cas9 (Clustered Regularly Interspaced Short Palindromic Repeats/Cas9) to create a homozygous isogenic control line (c.ID3).

##### *Human pluripotent stem cell cardiac differentiation*

Undifferentiated hiPSC cell lines were expanded under serum-free conditions in mTeSR medium (StemCell Technologies) on a Corning® Matrigel hESC-Qualified Matrix (StemCell Technologies), at 37°C, 5% $CO_2$. For cardiac differentiation, a monolayer directed differentiation protocol was applied using the cardiac PSC Cardiomyocyte Differentiation Kit (Life Technologies). HiPSC colonies at 80-85% confluence was chemically dissociated using 1X TrypLE (Life Technologies, Thermo Fisher Scientific, Carlsbad, CA, United States Life Technologies), suspended into mTeSR with 5 µM Y-27632 ROCK inhibitor (vendor), and seeded as single cells onto Matrigel-coated wells of a 24-well plate at a cell density of 80,000-100,000 cell/well. At ~85 % confluence, the medium was changed to Cardiomyocyte Differentiation Medium A (referred as day 0) to start cardiac induction. Medium A was replaced after 2 days with Medium B and Medium B was replaced after 2 days with Medium C for final differentiation. At ~ day 8-10 post-initiation of differentiation, the cells exhibited spontaneous beating. At this time, the differentiation medium was replaced with RPMI plus B-27 supplement (Life Technologies, Thermo Fisher Scientific, Carlsbad, CA, United States). Using this protocol, patient-derived cell lines (ID3), the relative isogenic control line (c.ID3) and a mEGFP-tagged α-actinin-2 hiPSC line (WTC Derived AICS hiPSC Lines, Allen Institute) were differentiated into cardiomyocytes. Specifically, hiPSC-derived cardiomyocytes expressing the mEGFP-tagged α-actinin-2 protein were dissociated into single cells and plated on nanopatterned surfaces (CuriBio 96-Well Plate* ANFS-0096) at a density of 10,000 cell/well to perform structural analyses.

##### *Engineered Heart Tissues generation*

Beating cardiomyocytes (day 15 p.d.) at ~90 % confluence were washed with 1X PBS and dissociated with 1X TrypLE for 10 min at 37 °C. The cell suspension was centrifuged at 1,000 rpm for 5 min and resuspended in cardiac medium with 10% FBS and 1 µM Y-27632 ROCK inhibitor. Fibrin EHTs were generated following previously described protocol (3,4)(21, 22). Briefly, agarose casting molds were prepared in 24-well tissue culture plates with 2 % Agarose (Life Technologies), dissolved in the appropriate volume of PBS; 1,6 mL of warm agarose was pipetted into the wells and then Teflon spacers (EHT Technologies GmbH) were inserted. After agarose solidification (~10 min), Teflon spacers were removed and silicone PDMS racks (EHT Technologies GmbH) were positioned upside down in the middle of each agarose well. EHTs consisted of 1 x $10^6$ cells in RPMI/B-27 with 5 mg/ml Fibrinogen from bovine plasma (Sigma-Aldrich) and 3 U/mL Thrombin (Sigma-Aldrich). After fibrin polymerization (37 °C) the silicone racks with attached fibrin gels were transferred to new 24-well plates and the medium was changed every two days using RPMI plus B-27 and 33 µg/ml Aprotinin (Sigma-Aldrich) (100 µL per EHT). After 8-10 days of culture, human EHTs showed spontaneous contractions, and deflection of the silicone pillars allowed video-optical analysis of the contraction.

##### *Contractile force measurements*

*Spontaneous contractions recordings:* Spontaneous auxotonic tension and beating frequency were regularly measured from day 20 to 50 of culture by optical tracking of the flexible post deflection at 10x magnification (Olympus) using an Evos FL2 auto system (20 seconds, 33 frame/sec) (Life Technologies) with an on-stage incubator connected to a BenchPro 2100 Plasmid Purification System (Invitrogen). The recordings were performed at 37°C, 5% $CO_2$ in the RPMI/B27 culture medium with ~0.4 mM of $Ca^{2+.}$ The relationship between the deflection of the individual pillars and force is 0.28 µN/ µM. Tension was analyzed using a customized Labview analysis program and EHT cross-sectional area (CSA) was calculated assuming an elliptical cross section (A= π/4 · width · thickness) for force normalization (5).

*Isometric force measurements:* For steady-state tension recording, EHTs were manually detached from the silicon pillars and immediately transferred into Krebs/Henseleit buffer, containing (in mM) 119 NaCl, 15 KCl, 0.5 $CaCl_2$, 1.2 $MgSO_4$, 1.2 $KH_2PO_4$, 25 $NaHCO_3$ with the addition of BDM (20 mM). Then the EHTs, like human trabeculae, were mounted between a force transducer (KG7A, Scientific Instruments Heidelberg, Germany) and a motor arm (Aurora Scientific Inc. Aurora, Canada), used to control the length of the tissue, and perfused with Krebs/Henseleit buffer, without BDM and 4.7 mM KCl, at pH 7.4 with 95%$O_2$:5%$CO_2$ and controlled by a custom Labview program (National Instruments, Austin, Texas). To normalize the force on the cross-sectional area (CSA), we calculated the thickness (c.ID3 0.68±0.04mm; ID3 0.69±0.05mm) and width (c.ID3 0.80±0.04mm; ID3 0.81±0.03mm) of each EHT, as they are not flat tissues. The width and thickness of the tissues were determined using a stereomicroscope with a reticle in the eyepiece and the thickness in particular was calculated thought the reflected image on a mirror placed at 45°. After fixation, the mechanical properties of EHTs were analyzed. The tissues were pre-stretched from $L_0$ (slack length) to $L_{max}$ (maximum force production length) according to Frank-Starling's law. The tissues were stretched with strain increments of approximately 3% up to the length at which the maximum contraction force was exerted and stimulated at 1Hz. After stabilization, isometric force was measured by stimulating the tissues at increasing pacing rates (0.2-2.5 Hz) and recordings were performed at 35 ± 2 °C at different $Ca^{2+}$ concentration (0.5, 1.8 and 4 mM). Isometric force was recorded via custom LabView software and was calculated using the same cross-sectional area normalization as for spontaneous measurements.

##### *Mavacamten long-term treatment*

For drug screening on EHTs and single hiPSC-CMs, Mavacamten (formerly MYK-461; MedchemExpress, USA) was dissolved in DMSO to obtain a 10 mM stock solution. For chronic treatment, Mavacamten was dissolved in DMSO at the dilution of 1mM and further diluted to 0.3 µM similarly to other studies (6,7) or 0.75 µM in the culture medium (RPMI/B-27). When the EHTs started to beat spontaneously, measurements were performed by optical tracking of the flexible post deflection before drug treatment for about 1 week. Then the EHTs were treated for 20 days with Mavacamten dissolved in RPMI/B-27. Spontaneous contractile force was monitored during the treatment. After the 20-day treatment, some EHTs were maintained in culture in the presence of the drug-free medium for 2 days and then tension recordings were performed under isometric conditions. The same protocol was applied to mEGFP-tagged alpha-actinin hiPSC cardiomyocytes, which were treated with Mavacamten (0.3 and 0.75 µM), 2,3-butanedione monoxime (BDM) (20 mM) and with phenylephrine (PE) (100 µM), from day 30 p.d. and monitored by EVOS FL2 auto microscope during cell culture. In parallel, single ID3 cardiomyocytes were also treated chronically with Mavacamten and Aficamten (0.3 and 0.75 µM). At the end of treatment, confocal microscope images were acquired.

##### *Image analysis*

The cell area and density of the sarcomere network were assessed with Image J software. Specifically, cell area was monitored during cell culture by analyzing bright-field images obtained with a fluorescence microscope, while analysis of sarcomere density was performed at the end of treatment on images acquired with a confocal microscope. In addition , the sarcomere network was determined by binary image generation, using the Image J ridge detection plugin, and filament density was calculated as a percentage of the total cell area (8).

##### *Action Potential recordings*

For dual recordings of APs and CaTs, EHTs (day 60 p.d.) were loaded with 2 µL/mL Fluovolt (Thermo Fisher, Waltham, MA, USA), 2 µL/ml of Cal630 (AAT Bioquest, Sunnyvale, CA, USA) and 5 µL of Power Load™ concentrate (Thermo Fisher, Waltham, MA, USA) for 30 min at 37 °C. The dyes were added to a Tyrode solution, containing (in mM) 5 HEPES, 10 Glucose, 140 NaCl, 5.4 KCl, 1.2 $MgCl_2$, 1.8 $CaCl_2$ (pH = 7.3) supplemented with 10 µM blebbistatin. After incubation, the EHTs were washed with pre-warmed culture medium before being placed in the experimental chamber. The experimental chamber features platinum electrodes for electrical field stimulation, connected to a stimulator (DigiTimer, Welwyn Garden City, UK). During measurements, tissues were continuously perfused with Tyrode's buffer containing blebbistatin to inhibit spontaneous contraction, and Mavacamten for EHTs that were chronically treated, and stimulated at different frequencies (1 Hz). For fluorescence studies, cells were simultaneously illuminated by LED light at two different wavelengths, blue (488 nm) for excitation of Fluovolt and yellow (580 nm) for Cal630 dye excitation, using a multi-led system (Lumencor SPECTRA X, Beaverton, OR, USA). A dual-wavelength band-pass filter cube (Semrock, IDEX, Lake Forest, IL, USA) was used to allow fluorescence light from the two dyes to be collected by a single camera (Photometrics Prime sCMOS, Teledyne, Tucson, AZ, USA).

##### *RNA isolation and transcriptome analysis*

Mutated and control EHTs untreated and treated with Mavacamten were also used to perform transcriptome analysis. RNA was isolated from each tissue and total transcriptome sequencing was performed to reveal gene expression levels. RNA isolation was carried out using QIAzol Reagent to facilitate tissue lysis and the RNeasy Kits for RNA Purification (© QIAGEN, Hilden, Germany). Quantification of the isolated RNA was performed using the NanoDrop™ spectrophotometer (Thermo Fisher Scientific, Carlsbad, CA, United States). RNA sequencing (RNA-seq) to reveal gene expression levels was performed by Breda Genetics (Brescia, Italy). Raw sequencing reads were quantified using Salmon (v1.10.1) with the parameters --seqBias, --gcBias, and --validateMapping, aligned against the human reference transcriptome (GRCh38) (9). Downstream analysis was conducted in R (v4.2.1) using the DESeq2 package (v1.37.6) (10). Expression differences were assessed across three experimental comparisons: mutated untreated vs. isogenic untreated, mutated treated with 0.3 µM Mavacamten vs. mutated untreated, and mutated treated with 0.75 µM Mavacamten vs. mutated untreated. Quality control (QC) based on Principal Component Analysis (PCA) and sample distance metrics revealed the presence of low-quality samples, which were excluded from the following steps. For the first comparison, samples originating from two different batches were corrected for batch effects using the ComBat algorithm implemented in the sva package (v3.46.0) (11). For the second and third comparisons, which involved three experimental batches, technical variation was mitigated using the Remove Unwanted Variation (RUV) approach from the RUVSeq package (v1.32.0) (12), providing a more robust correction across conditions. Similarly, for the third comparison, batch correction was applied. Across all comparisons, differentially expressed genes (DEGs) were identified using thresholds of adjusted p-value < 0.05 and absolute $\log_2$ fold change > 0.5. Gene set enrichment analysis (GSEA) was performed for each set of DEGs using clusterProfiler (v4.6.2) (13), referencing the Gene Ontology (GO) and KEGG pathway databases to identify significantly enriched biological processes and pathways related to hypertrophic cardiomyopathy. For KEGG enrichment in the second comparison, a more stringent filter (adjusted p-value < 0.01) was applied.

##### *Statistical analysis*

All data are expressed as mean ± SEM and the number of hiPSC-CM differentiations (N) and the number of EHTs or individual cells (n) are indicated in the respective legends. Normality and homogeneity of variance were verified before applying parametric tests. Comparisons were made using a two-tailed Student's *t* test and differences between groups were considered statistically significant when $*p \leq 0.05$. One-way ANOVA with a Tukey post-hoc test was used to compare multiple groups. Probability levels considered as statistically significant were * $p < 0.05$, ** $p < 0.01$, *** $p<0.0001$. Graph analysis was performed using OriginPro software (OriginLab, Northampton, Massachusetts, USA).

### SUPPLEMENTARY FIGURES

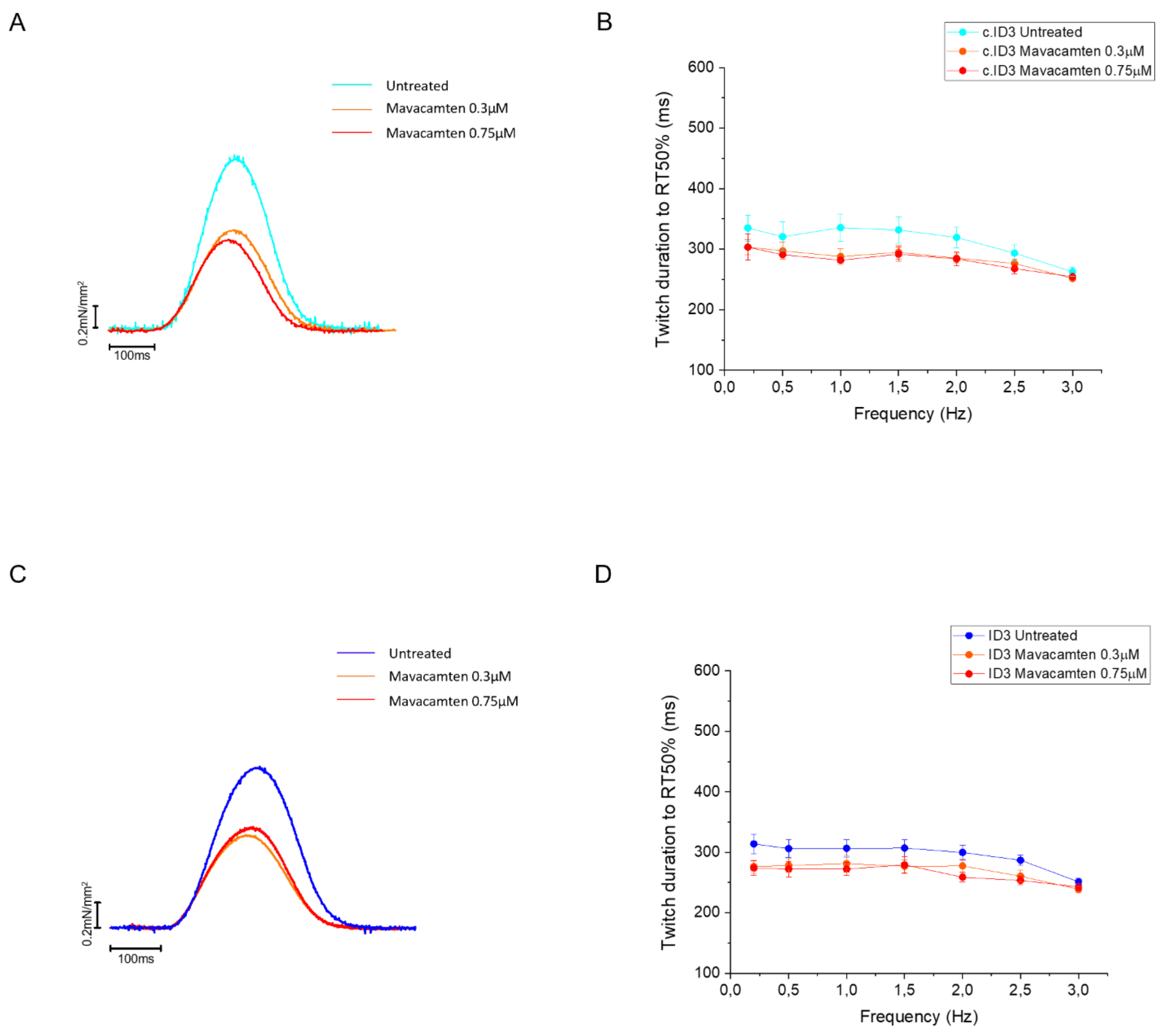


**Figure S1. Twitch duration of EHTs at different pacing frequencies after long-term Mavacamten exposure and after 4 days of washout.** (A-C) Representative twitches of untreated and Mava-treated EHTs. (B-D) Twitch duration at 50% of relaxation at 0.2-2.5 Hz of EHTs treated and untreated with Mavacamten. EHTs were measured in isometric conditions at 37°C in the Krebs-Henselheit solution with 1.8 mM of [$Ca^{2+}$] under imposed pacing. Error bars ± SEMs. Statistical analysis was performed using one-way ANOVA with Tukey post-hoc test. No statistically significant differences emerged. (c.ID3, n=5; ID3, n=6).

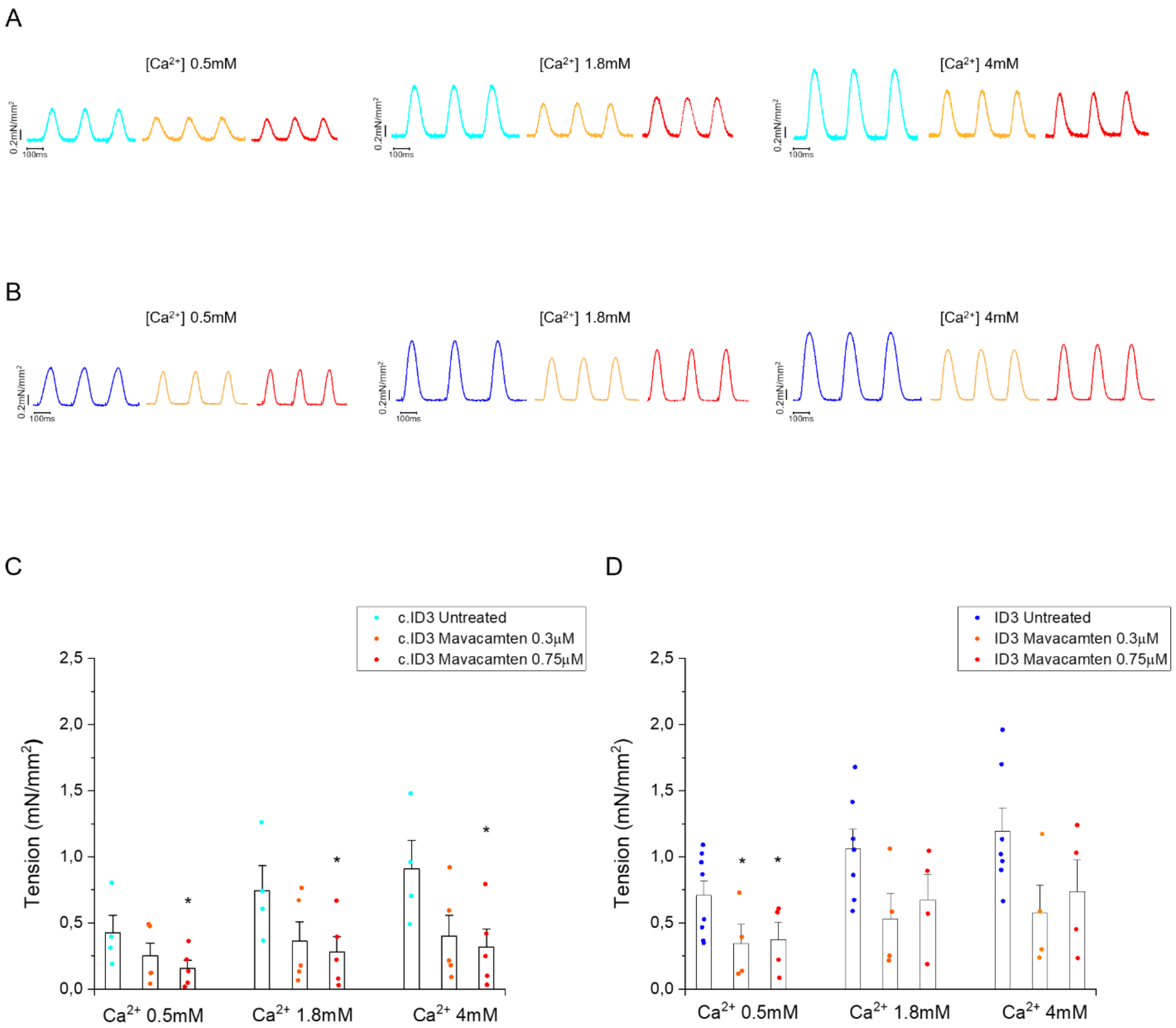


**Figure S2. Inotropic response of c.ID3- and ID3- EHT after chronic treatment and 4 days washout.** Representative twitches of untreated and Mavacamten-treated c.ID3- (A) and ID3-EHTs (B) at different extracellular [$Ca^{2+}$]. (C-D) Active tension at 1 Hz constant pacing of c.ID3- and ID3-EHTs treated with Mavacamten 0.3 µM and 0.75 µM compared with untreated tissues. EHTs were measured in isometric conditions at 37°C in the Krebs-Henselheit solution with 0.5 mM,1.8 mM and 4 mM of [$Ca^{2+}$] under imposed pacing. Error bars ± SEMs. Statistical analysis was performed using one-way ANOVA with Tukey post-hoc test. No statistically significant differences emerged.

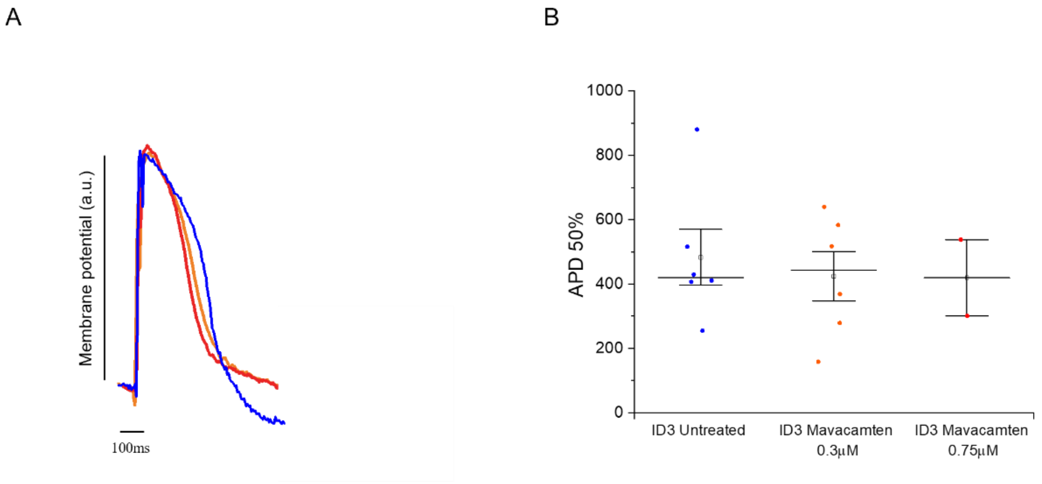


**Figure S3. Action potential recording by fluorescent indicator (FluoVolt).** Untreated ID3-EHTs (N=5, n=6) were compared with ID3-EHTs treated with Mavacamten at 0.3 μM (N=7, n=6) and 0.75 μM (N=2, n=2) for action potential at 50% of duration (APD50, ms), stimulated at 0.5Hz. Error bars ± SEMs.

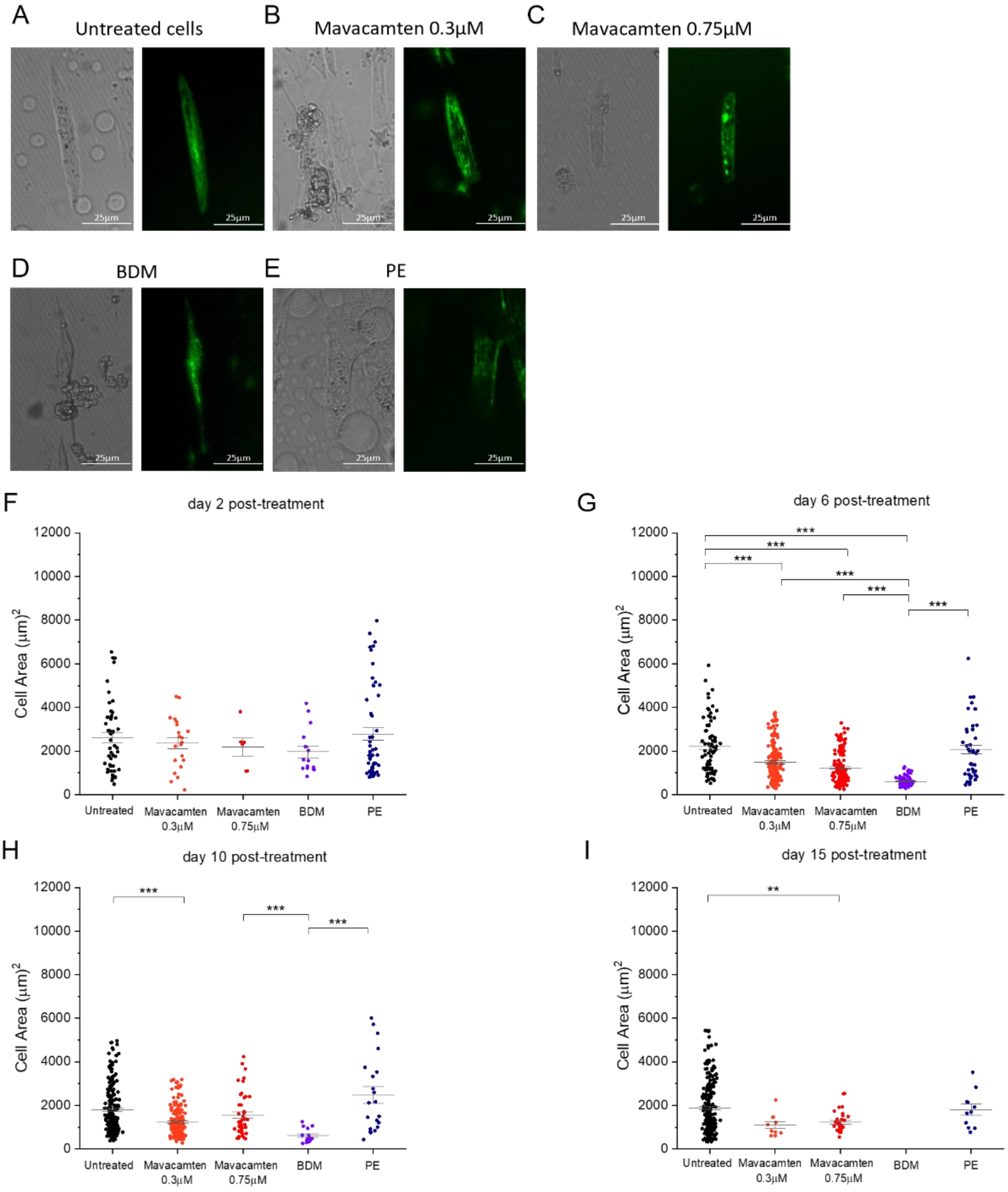


**Figure S4. Time point analysis of cell area in cardiomyocytes derived from single mEGFP-tagged α-actinin hiPSC.** (A-E) Bright-field and fluorescence microscope images of mEGFP-tagged α-actinin cells during the treatment. (F-G-H-I) Analysis of cell area during cell maturation in the presence of Mavacamten (0.3 and 0.75 µM), BDM (20 mM) and PE (100 µM). Statistical analysis was performed using one-way ANOVA, with a Tukey post-hoc test with statistical significance set at * $p < 0.05$, ** $p < 0.01$, *** $p<0.0001$.

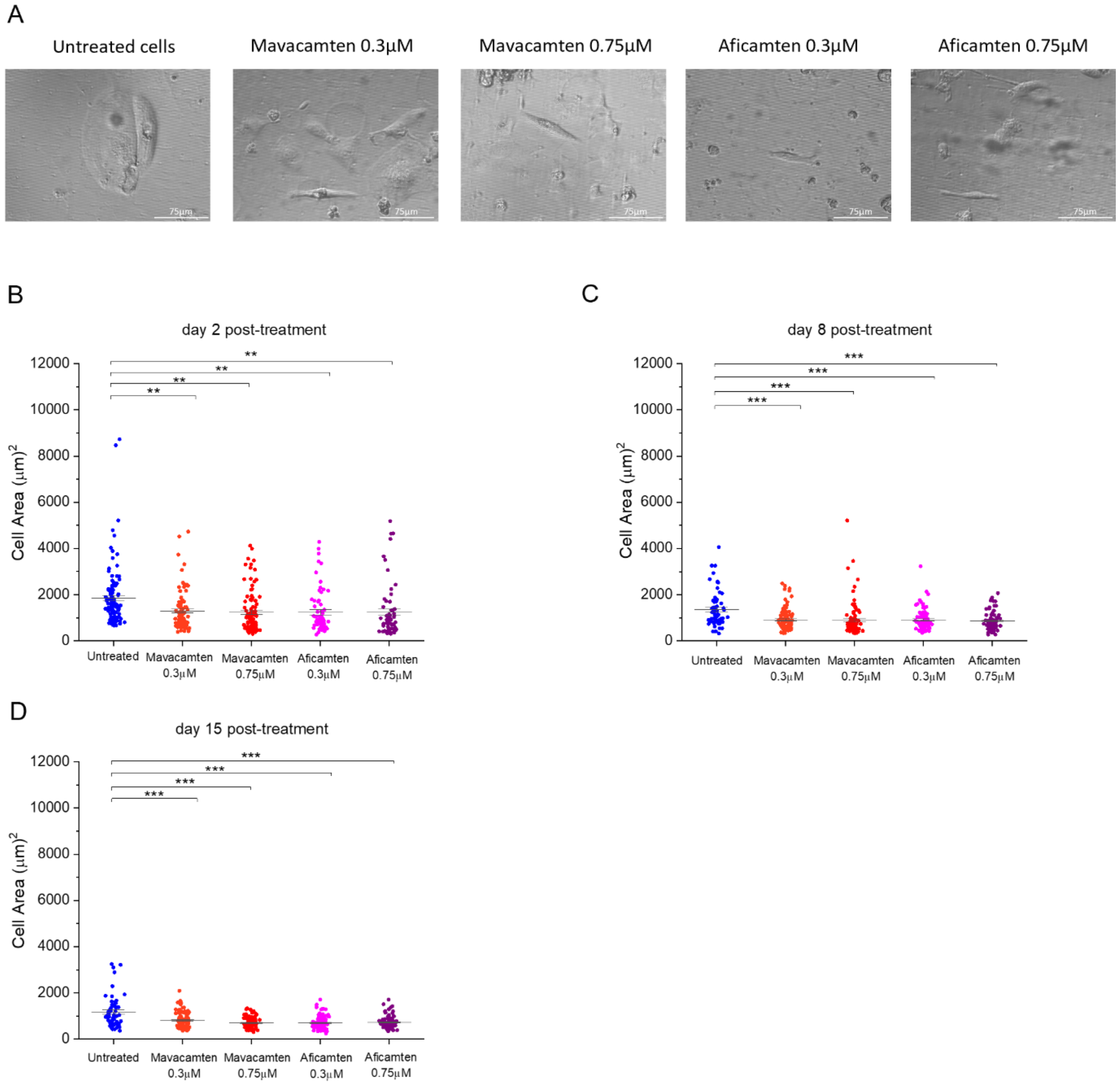


**Figure S5. Time point analysis of cell area in cardiomyocytes derived from ID3 hiPSC.** (A) Bright-field images of c.772G>A hiPSC-CMs in the presence of Mavacamten (0.3 and 0.75 µM) and Aficamten (0.3 and 0.75 µM). (B-D) Analysis of cell area during cell maturation and the treatment with Mavacamten and Aficamten. Error bars ± SEMs. Statistical analysis was performed using one-way ANOVA, with a Tukey post-hoc test with statistical significance set at * $p < 0.05$, ** $p < 0.01$, *** $p<0.0001$.

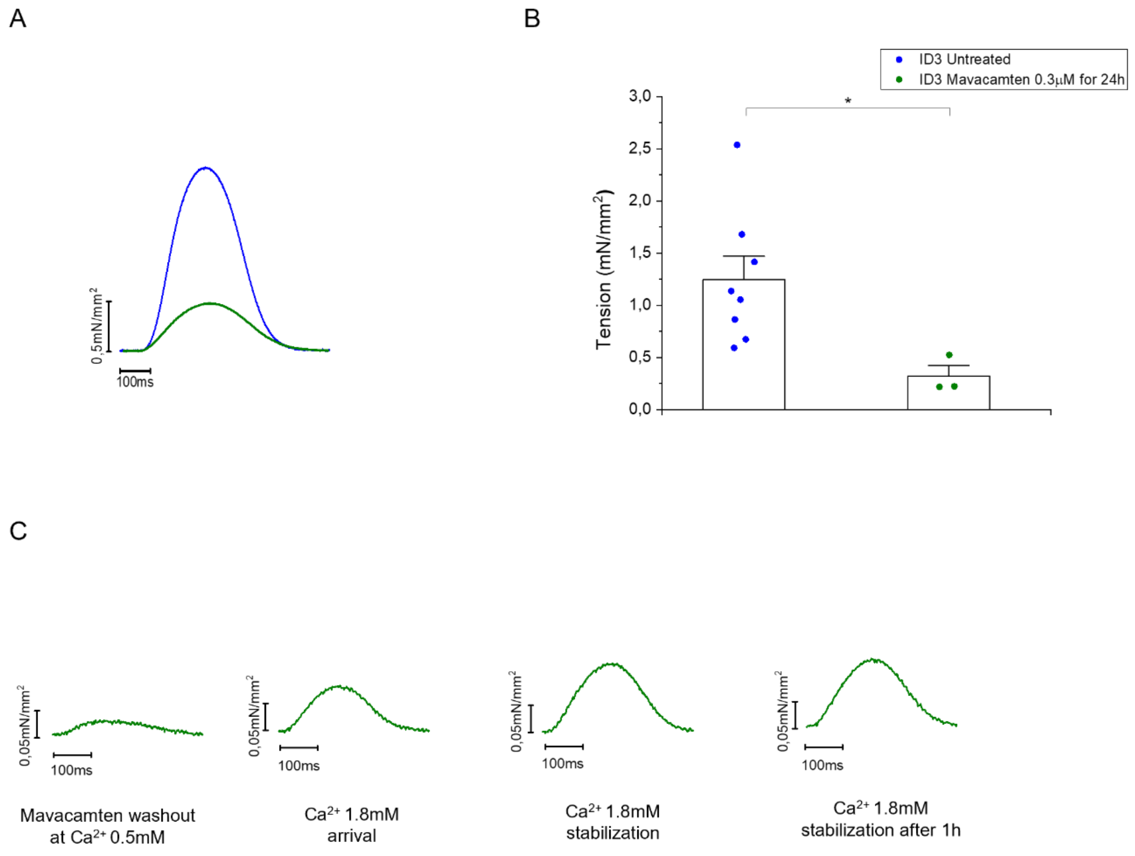


**Figure S6. Contractile properties of EHTs at advanced stages of maturation following 24 hours with Mavacamten 0.3µM.** (A-B) Active tension of ID3-EHTs (N=5; n=8) compared to ID3-EHTs treated with Mavacamten 0.3 µM for 24 hours at day 55 p.d. (N=1; n=3). Recordings were performed at 1 Hz constant pacing with 1.8 mM of [$Ca^{2+}$]. Error bars ± SEMs. Statistical analysis was performed using one-way ANOVA, with a Tukey post-hoc test with statistical significance set at * $p < 0.05$.

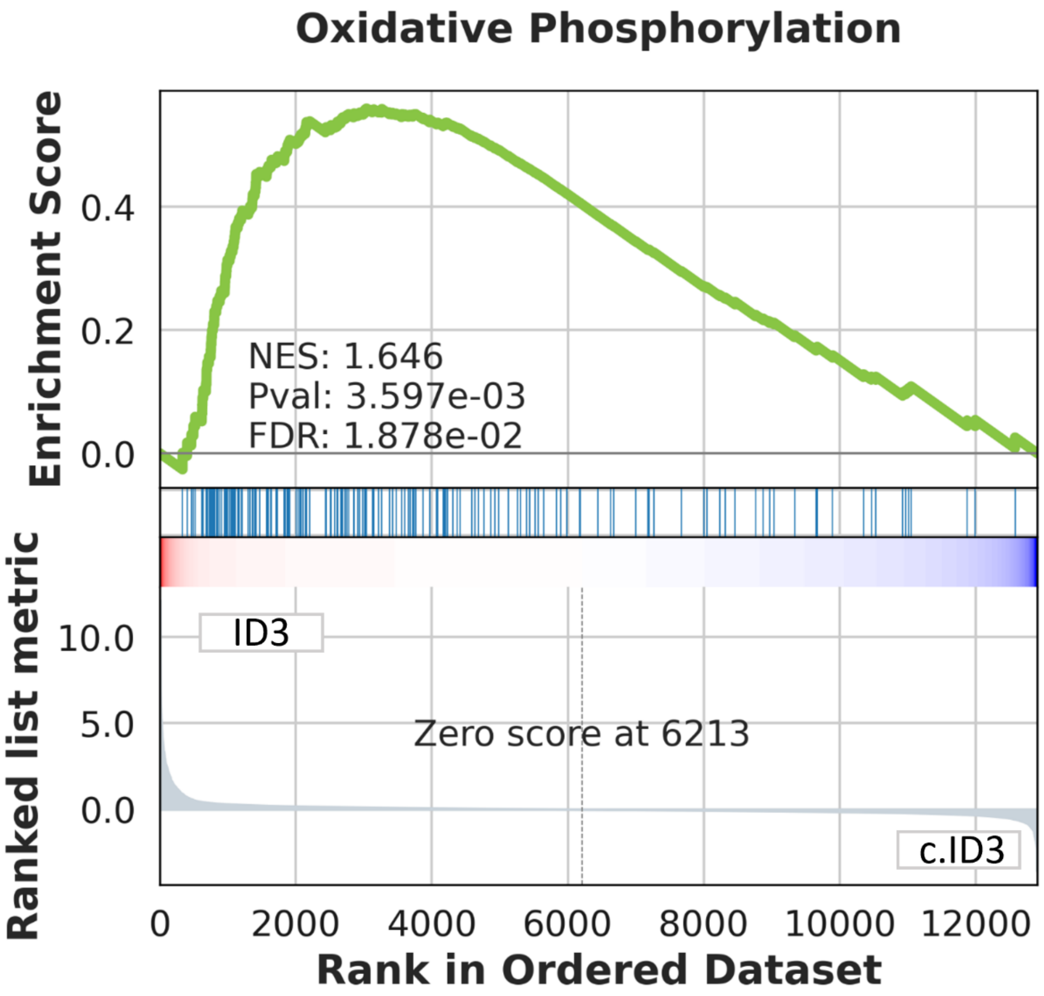


**Figure S7. Gene Set Enrichment Analysis (GSEA) plot for the Oxidative Phosphorylation pathway comparing untreated ID3 versus c.ID3 samples**. The enrichment score (green line) reflects the degree to which genes in the oxidative phosphorylation pathway are overrepresented at the top or bottom of the ranked list of genes. The maximum enrichment score (NES) is 1.646, with a nominal p-value of 0.0036 and an FDR (false discovery rate) of 0.0188, indicating significant enrichment in the ID3 condition. The vertical blue lines in the middle panel indicate the position of pathway genes in the ranked gene list. The enrichment is skewed towards the top of the ranked list (ID3), suggesting upregulation of oxidative phosphorylation genes in HCM samples (n=4).

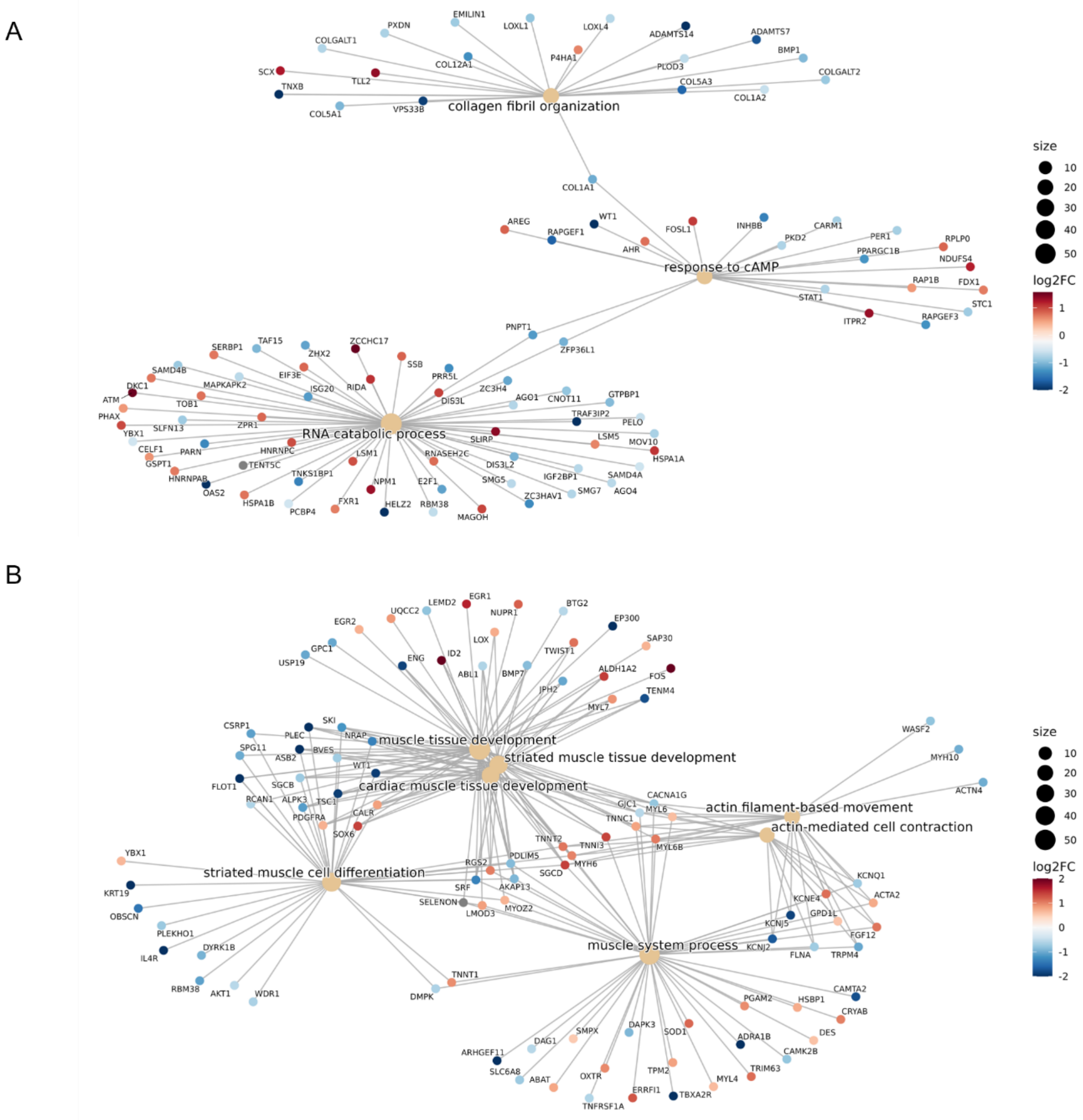


**Figure S8. Exclusive Gene Ontology enrichment network of Mavacamten treated EHTs.** (A) Gene Ontology (GO) enrichment network of differentially expressed genes (DEGs) for the sample treated with 0.3 µM Mavacamten (n=3). GO terms (yellow nodes), corresponding to those identified in the lower panel of Figure 8, are connected to their associated DEGs (coloured nodes), filtered by statistical significance (adjusted p-value < 0.05). Node colour represents the log2 fold change (log2FC) in gene expression, and node size reflects the number of genes associated with each GO term.(B) GO enrichment network of DEGs for the sample treated with 0.75 µM Mavacamten (n=3). As in panel A, grey nodes represent GO terms previously identified in Figure 8, while coloured nodes indicate significant DEGs (adjusted p-value < 0.05). Node colour and size respectively indicate log2FC and gene count per GO term.

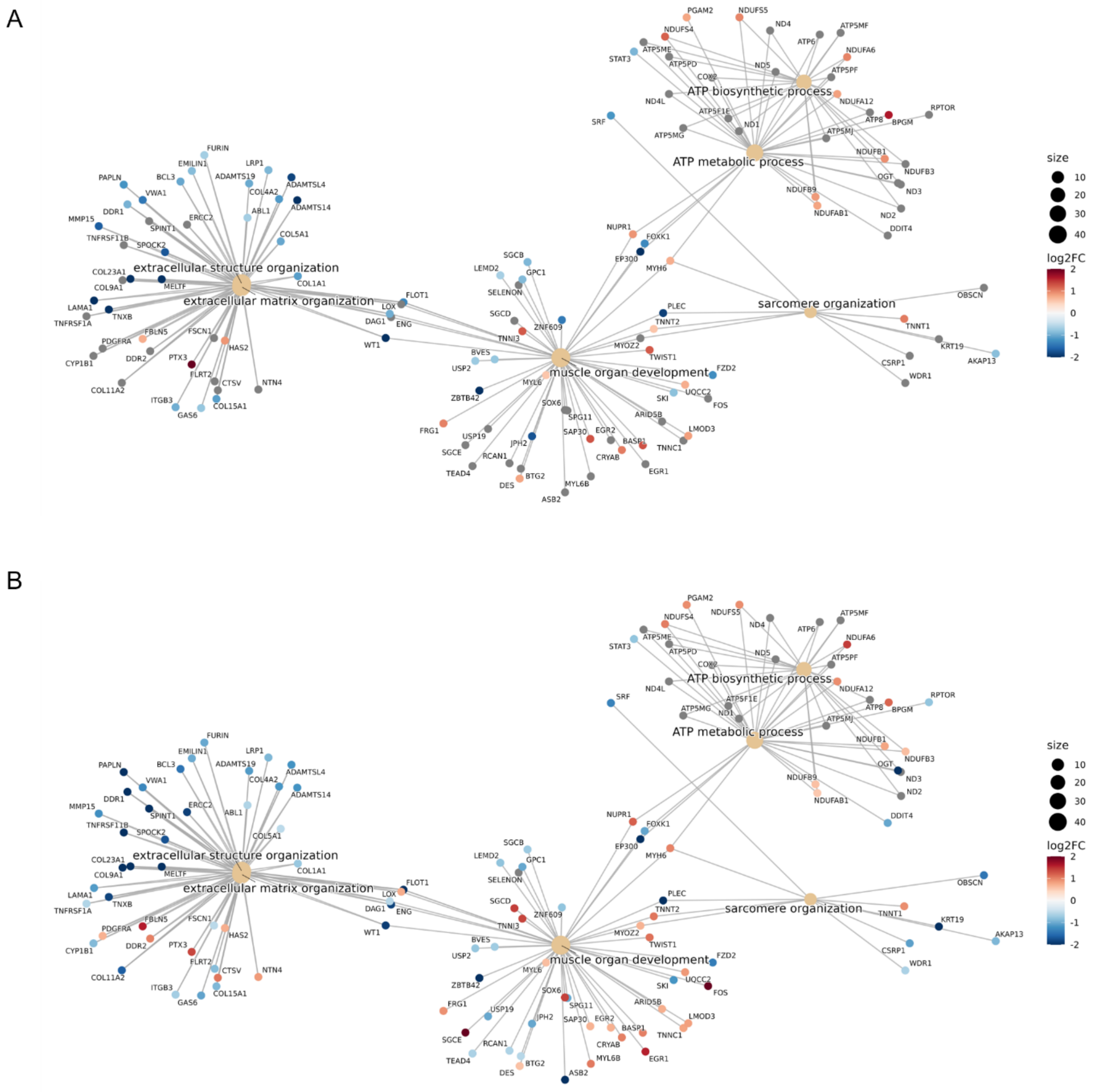


**Figure S9. Common Gene Ontology enrichment network of Mavacamten treated EHTs.** (A) Gene Ontology (GO) enrichment network of differentially expressed genes (DEGs) for the sample treated with 0.3 µM Mavacamten (n=3). GO terms (yellow nodes), corresponding to those identified in the upper panel of Figure 8, are connected to their associated DEGs (coloured nodes), filtered by statistical significance (adjusted p-value < 0.05). Node colour represents the log2 fold change (log2FC) in gene expression, and node size reflects the number of genes associated with each GO term. (B) GO enrichment network of DEGs for the sample treated with 0.75 µM Mavacamten (n=3). As in panel A, yellow nodes represent GO terms previously identified in Figure 8, while coloured nodes indicate significant DEGs (adjusted p-value < 0.05). Node colour and size respectively indicate log2FC and gene count per GO term.

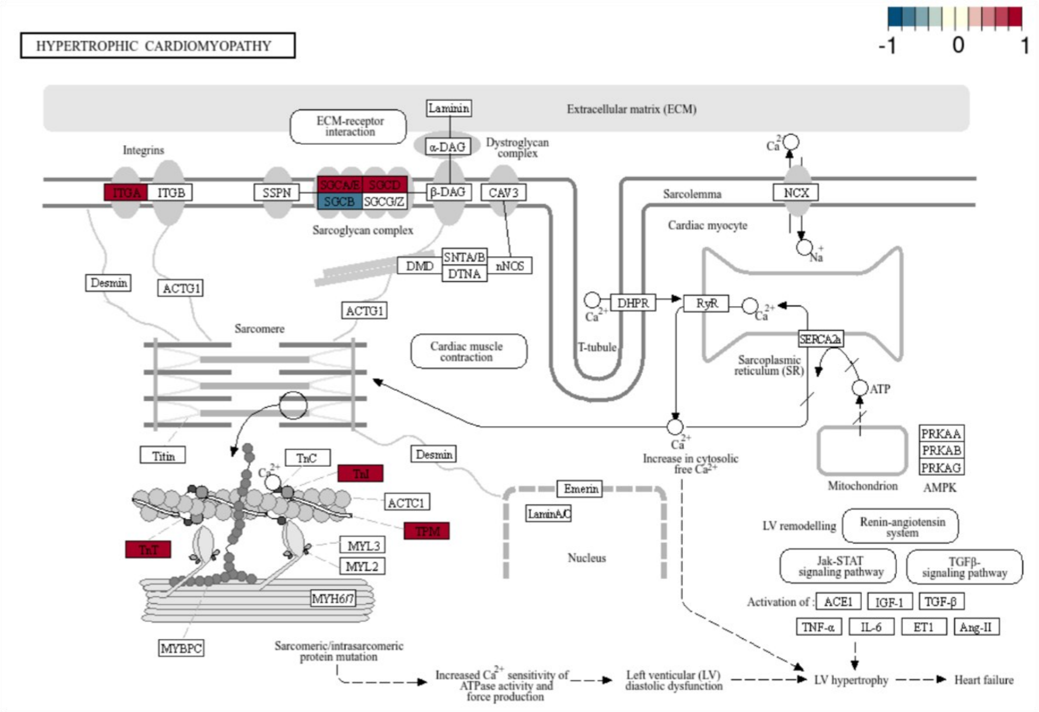


**Figure S10. KEGG pathway map of Hypertrophic Cardiomyopathy highlighting gene expression changes in the sample treated with 0.75 µM Mavacamten.**

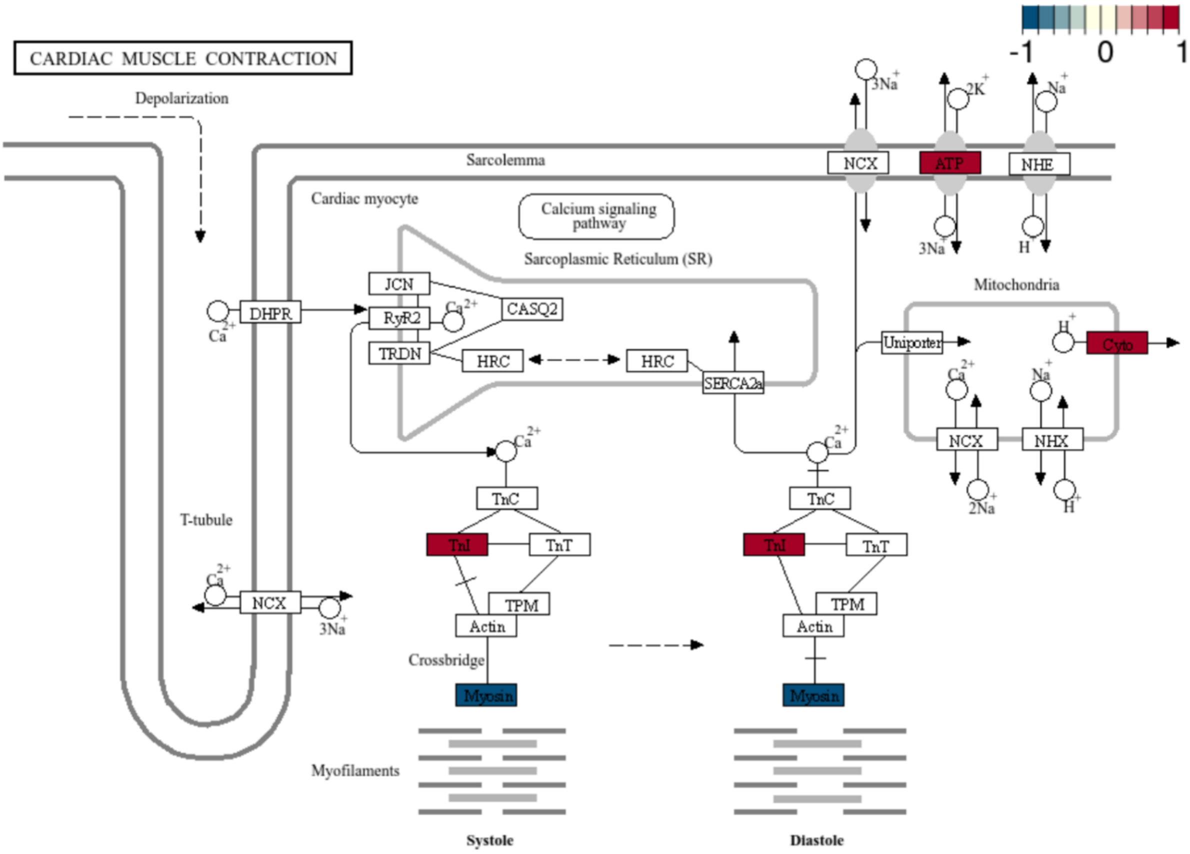


**Figure S11. KEGG pathway map of Cardiac Muscle Contraction highlighting gene expression changes in the sample treated with 0.3 µM Mavacamten.**

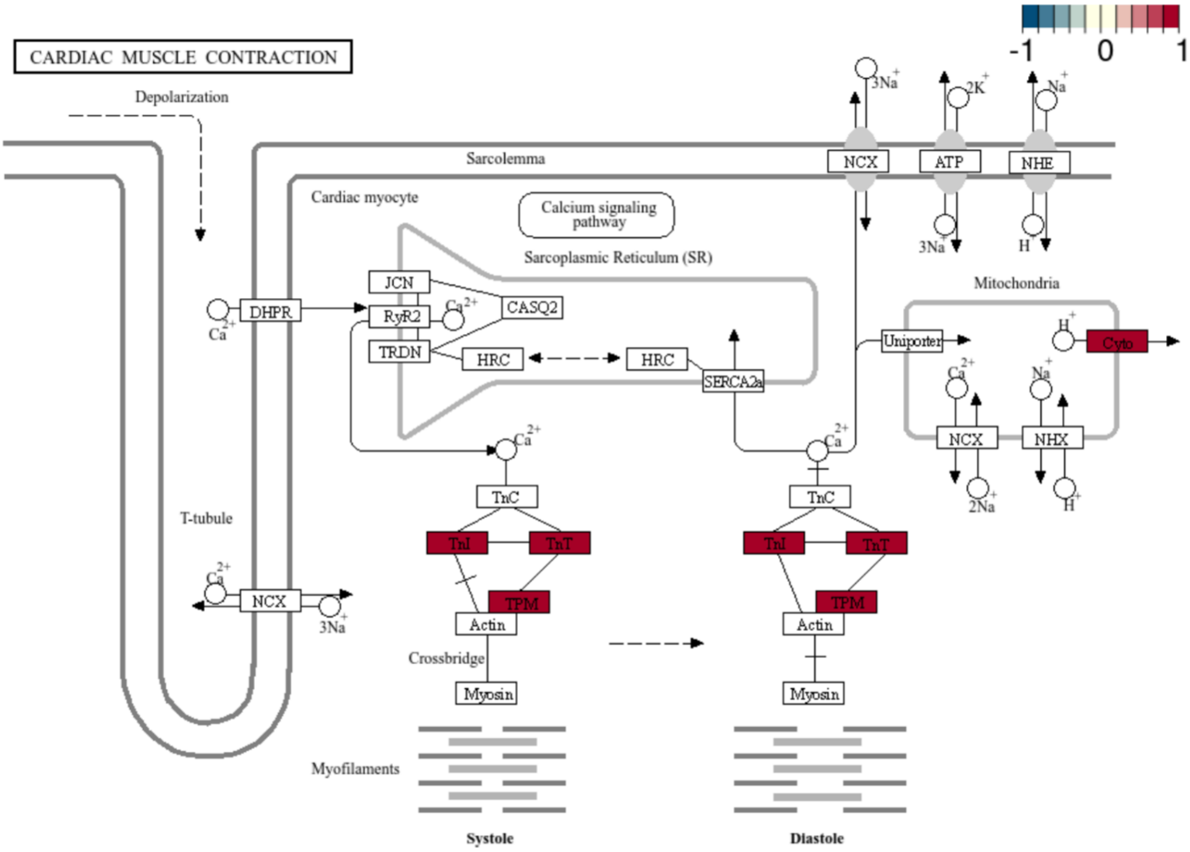


**Figure S12. KEGG pathway map of Cardiac Muscle Contraction highlighting gene expression changes in the sample treated with 0.75 µM Mavacamten.**

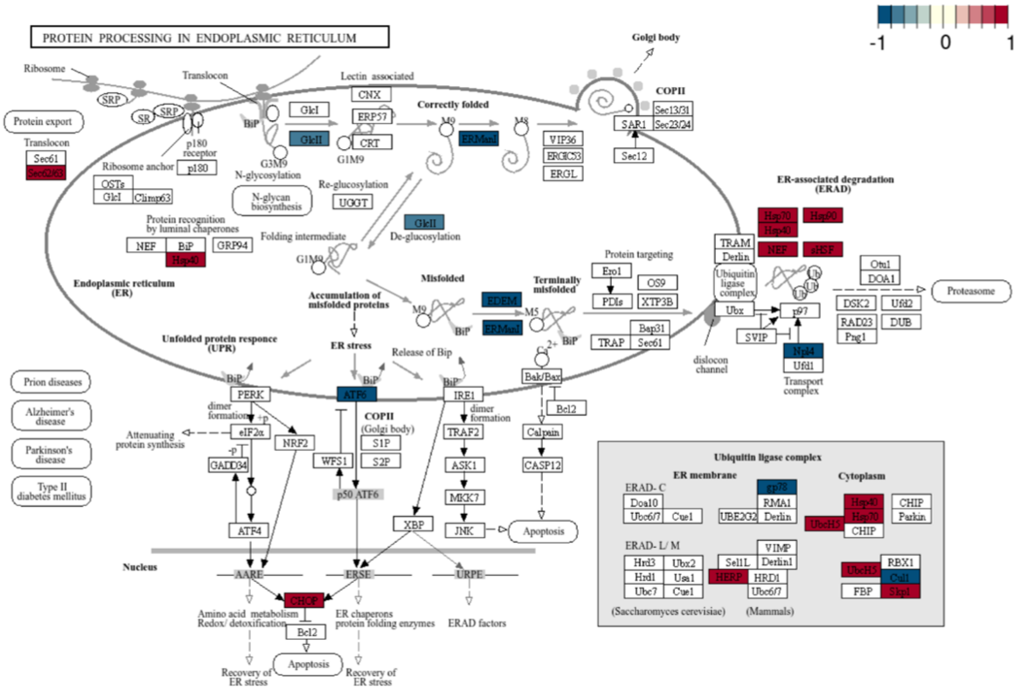


**Figure S13. KEGG pathway map of Protein Processing in Endoplasmic Reticulum highlighting gene expression changes in the sample treated with 0.3 µM Mavacamten.**
